# Distinct Narrow and Broadband Gamma Responses in Human Visual Cortex

**DOI:** 10.1101/572313

**Authors:** Eleonora Bartoli, William Bosking, Ye Li, Michael S. Beauchamp, Daniel Yoshor, Brett L. Foster

## Abstract

High frequency activity (> 30 Hz) in the neocortical local field potential, typically referred to as the ‘gamma’ range, is thought to have a critical role in visual perception and cognition more broadly. Historically, animal studies recording from visual cortex documented clear narrowband gamma oscillations (NBG; ∼20-60 Hz) in response to visual stimuli. However, invasive measurements from human neocortex have highlighted a different broadband or ‘high’ gamma response (BBG; ∼70-150+ Hz). Growing evidence suggests these two forms of gamma response are distinct, but often conceptually or analytically conflated as the same ‘gamma’ response. Furthermore, recent debate has highlighted that both the occurrence and spectral properties of gamma band activity in visual cortex appears to be dependent on the attributes and class of presented visual stimuli. Using high-density intracranial recordings from human visual cortex, we integrate and extend these findings, dissociating the spectral, temporal and functional properties of NBG and BBG activity. We report results from two experiments, manipulating visual stimulus attributes (contrast-varying gratings) and class (object categories) dissecting the differential properties of NBG and BBG responses. NBG oscillations were only reliably recorded for grating stimuli, while their peak frequency varied with contrast level. Whereas BBG activity was observed in response to all stimulus classes tested, with no systematic change in its spectral features. Temporally, induced NBG was sustained throughout stimulus presentation, in opposition to a more transient response for the BBG. These findings challenge the ubiquity of ‘gamma’ activity in visual cortex, by clearly dissociating oscillatory and broadband effects.

**Significance Statement:** Neocortical narrowband gamma oscillations (∼20-60 Hz) have been implicated in vision and cognition as a mechanism for synchronizing brain regions. Efforts to study this phenomenon have revealed an additional ‘high-gamma’ range response (∼70-150+ Hz), which is broadband and non-oscillatory. These different gamma range activities are often conflated in support of the same functional role. Using invasive recordings from human visual cortex, we show that narrow and broadband gamma can be dissociated by spectral, temporal and functional response properties. While broadband gamma responses were more transient to the presentation of all stimuli, narrowband gamma responses were sustained and only occurred reliably to grating stimuli. These differences have important implications for the study, analysis and interpretation of neocortical gamma range activity.

## Introduction

Neocortical gamma oscillations, rhythmic neural population activity in the ∼20-60 Hz frequency range, have historically been implicated in visual perception and cognitive processing more generally (1-4). Early theoretical accounts posited that the synchronization of gamma oscillations within visual cortex supported the binding of disparate visual features (3-5). This view has since been extended to propose the synchronization of gamma oscillations as a more general mechanism for inter-areal communication in neocortex, supporting higher cognitive function (2). Central to these theories is the capacity for gamma oscillations to provide rhythmic control over spiking activity, allowing temporally coincident spiking between synchronized neocortical areas (6).

After early studies identifying neocortical gamma oscillations in the cat and non-human primate (7-9), investigations of the human brain sought to identify similar gamma range activity patterns. While early non-invasive methods, such as electroencephalography (EEG), provided evidence of neocortical gamma oscillations (10), the >30 Hz frequency range presented challenges in dissociating small amplitude high-frequency activity from biological noise occupying a similar frequency range (11, 12). Subsequent work using human intracranial recordings, which have superior sensitivity to high frequency activity, suggested clear evidence of gamma range (>30 Hz) responses (13). However, in contrast to earlier animal work, these human studies most reliably identified activity in a higher and broader frequency range (e.g. spanning from 70 up to 150-200 Hz) leading many to describe these response patterns as ‘high-gamma’ (13). Importantly, this high-gamma range activity has been repeatedly observed across neocortical regions beyond visual cortex, and is now commonly used as an indicator of local electrocortical response (14).

Despite these apparent spectral differences in what we will term narrowband gamma (NBG, i.e. gamma oscillations) and broadband gamma (BBG, i.e. high-gamma), these two signals are often conceptually and analytically conflated. However, growing evidence suggests these spectrally distinct responses are also temporally dissociated owing to different biophysical generators (15). Whereas NBG appears oscillatory, reflecting synchronization in the local field potential (LFP), BBG is often observed as non-oscillatory, potentially reflecting local population spiking and other activities coincident with multi-unit activity (MUA) (15-17). NBG and BBG may also be functionally dissociated, as growing evidence suggests NBG, unlike BBG, is highly stimulus dependent in at least two ways. Firstly, properties such as the amplitude and frequency of NBG are dependent on stimulus attributes (attribute dependence; e.g. stimulus size or contrast) (18-21). Secondly, the occurrence of NBG itself may be contingent on specific types of stimuli, showing a strong preference for commonly used visual gratings, rather than complex or natural stimuli (class dependence) (22).

Gamma range activity is readily used as a signature for sensory or cognitive processing in many areas of systems neuroscience (2). Therefore, the issue of clearly dissociating NBG and BBG activities to elucidate their spectral, temporal and functional differences is critical in adjudicating the genuine role these activates play in sensory and cognitive processing. However, sensitive measurement of these responses can be challenging using non-invasive techniques and invasive studies have predominantly been performed in non-human primates, with some key exceptions (21, 22). It is, therefore, important to integrate these findings across species by using similar experimental manipulations and comparable measurement resolution in larger study cohorts. To achieve this, we used high-density macro- and mini-ECoG recordings from human visual cortex during two visual experiments that aimed to induce, manipulate and dissociate NBG and BBG. Overall, we observed a sustained amplitude increase within a narrow frequency range (i.e. NBG) to be induced reliably only by grating stimuli, whose contrast level determined the peak frequency of the induced oscillation. This stimulus dependence was not observed at higher frequency ranges, which showed a broadband increase in amplitude without any characteristic spectral peak (i.e. BBG), occurring transiently at stimulus onset and offset and similarly across different visual stimuli. Our findings have clear import for the detection, interpretation and functional role of neocortical gamma range activity in vision and cognition more broadly.

## Results

### Identification of responsive sites in early visual cortex

In the current study, we used high-density electrocorticography (ECoG) recordings from human visual cortex in 7 subjects undergoing invasive monitoring for the surgical treatment of refractory epilepsy (see Methods). Recordings were performed using hybrid electrodes arrays, where mini-ECoG electrodes (0.5 mm diameter) were fabricated in-between standard macro-ECoG electrode (2 or 3 mm diameter) strip arrays in two configurations (see Figure 1A and Figure S1). We employed a functional criterion to identify responsive electrodes within early visual cortex based on the presence of a visual-evoked potential (VEP; see Methods). The VEP served as a measure of visual responsiveness that is partially independent from NBG and BBG. Of all the electrodes anatomically localized to the occipital lobe, approximately 65% (133/205 electrodes) displayed a VEP, with evoked components similar to those classically identified in scalp (e.g. N1 occurring around ∼70 ms) and intracranial recordings (23). VEPs are depicted for all electrodes in Figure 1B, while the anatomical location of macro/mini ECoG electrodes with/without a VEP is represented in Figure 1C on a common cortical surface. Subsequently, electrodes not displaying a VEP were discarded from further analysis, as the absence of such a response implied that the cortical location was either not in early visual cortex or that the electrode itself had poor sensitivity (e.g., not making good contact with the cortical surface, due to the presence of large veins or a slight torsion on the electrode array). For each selected electrode, we next analyzed the spectral response to visual grating stimuli.

**Figure 1.**
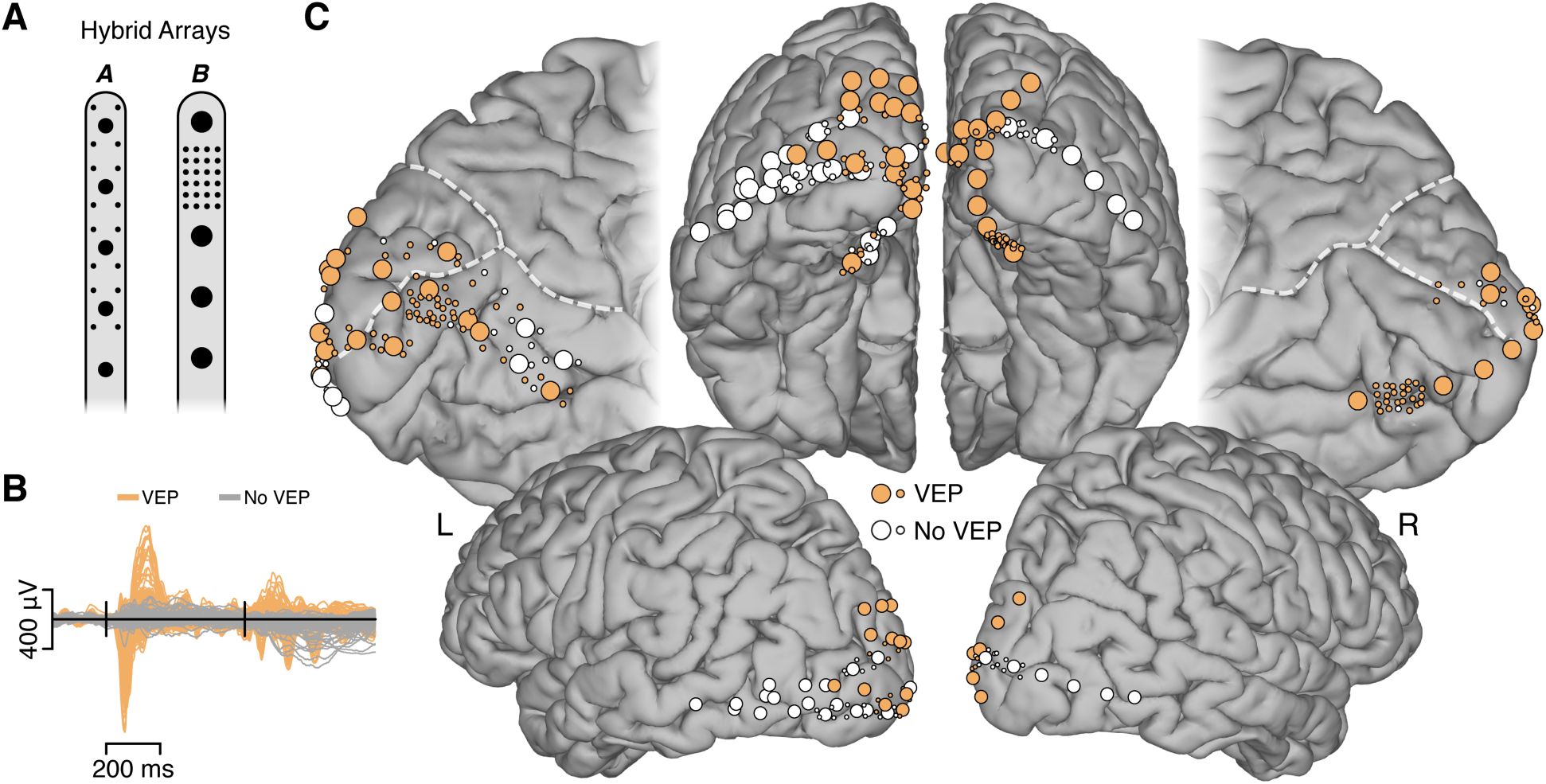
Electrode arrays and visually responsive recording sites. **A)** Schematic of hybrid macro-/mini-ECoG electrode arrays employed in this study. Standard clinical strip arrays with macro electrodes were customized to include small diameter mini electrodes in two configurations (array A = subjects 1-5; array B = subjects 6-7). Macro electrodes had diameters of 2 mm (array A) and 3 mm (array B), with mini electrodes having a diameter of 0.5mm (array A & B). **B)** To functionally select visually responsive electrodes, we used the visual evoked potential (VEP; see Methods). Mean voltage traces are shown for VEP (orange) and non-VEP (gray) electrodes. Black vertical lines indicate stimulus onset and offset. From a total of 205 electrodes, 133 (∼65%) displayed a VEP. **C)** Anatomical location of electrodes from all subjects are shown on a standard cortical surface (see Methods and Figure S1 for single subject locations). Electrodes were located on the medial, lateral and posterior aspects of the occipital lobe. Electrodes displaying a VEP are shown in orange (non-VEP in white). Dashed white lines indicate the parieto-occipital and calcarine sulci.

### Visual grating stimuli induce NBG and BBG responses

In experiment 1, our first aim was to quantify gamma range activity in early visual cortex in response to visual gratings. Subjects were presented with full screen static grayscale grating stimuli (spatial frequency 1 cycle/degree) for 500 ms. Gratings were presented at three contrast levels (20%, 50% and 100%), and two orientations (0° and 90°). Subjects were required to maintain fixation and respond to an oddball target grating (at 45° orientation, discarded from further analyses) by pressing a button. For many recordings, grating stimuli induced clear oscillatory responses in the raw ECoG voltage, identifiable on single trials, as exemplified in Figure 2A. Time-frequency analysis revealed these oscillatory responses to be induced NBG activity. Group averaged time-frequency plots (Figure 2B) indicate that induced NBG shows a sustained increase in amplitude, within a frequency range (∼ 20-60 Hz) consistent with prior human and non-human primate studies (24). In addition, at stimulus onset and offset transient amplitude increases were observed which extend into a higher BBG (∼ 70-150 Hz) range. These group spectrograms include both macro- and mini-ECoG electrodes, as both electrode types showed highly consistent responses and are therefore combined in subsequent analysis (see Figure S2 and S6). Next, we examined how these gamma range responses were influenced by changes in visual grating contrast.

**Figure 2.**
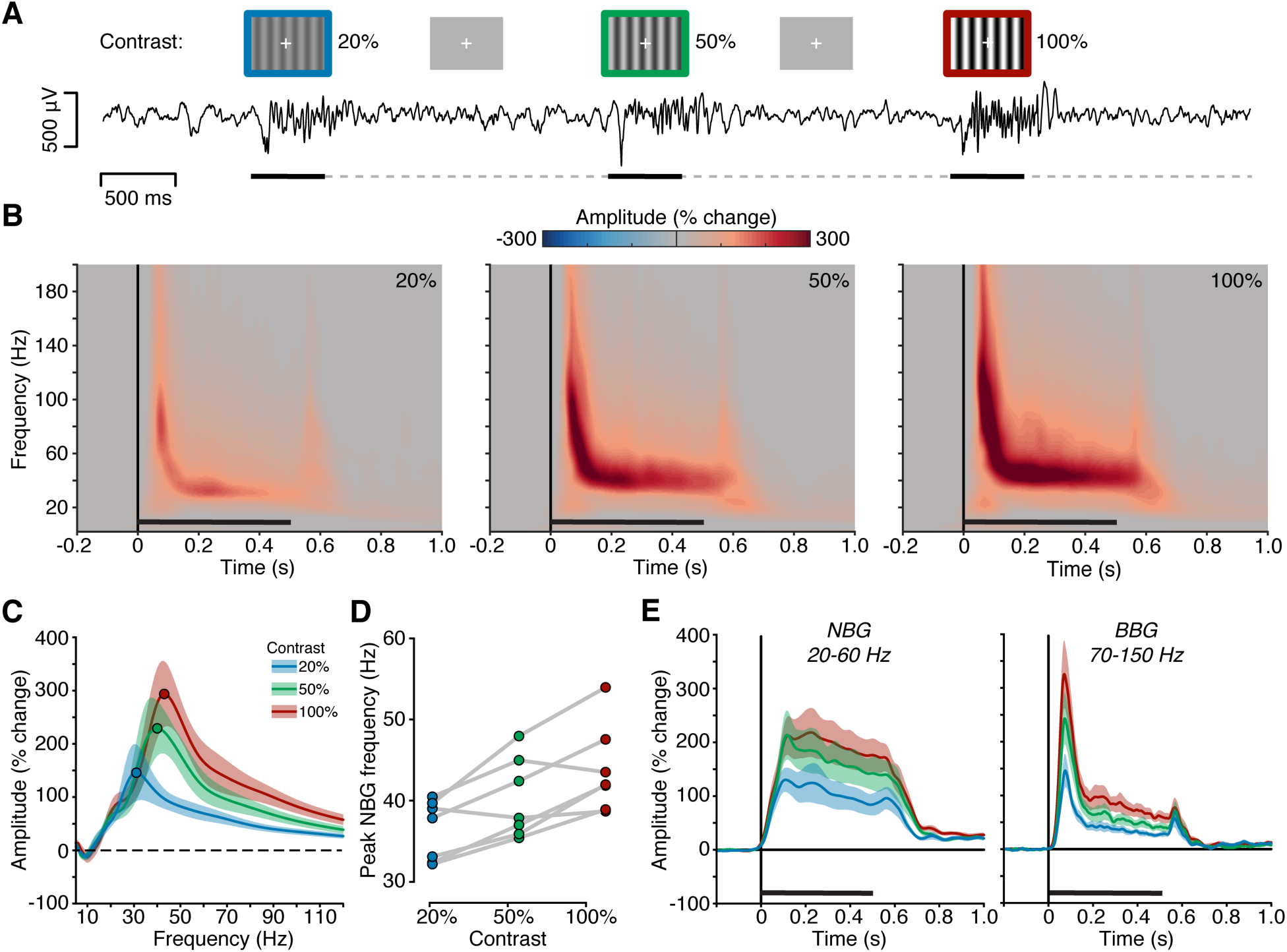
Experiment 1, spectral response to visual grating stimuli. **A)**. Experiment 1 stimuli and example voltage response to stimuli is shown (from subject N6). Static grating stimuli were presented for 500 ms (black line), with a random inter-stimulus interval (ISI; dashed line) of 1.5-2.0 s (see Methods). Gratings were presented at 20%, 50% and 100% contrast levels (blue, green and red borders reflect condition labels respectively and were not shown during the experiment). Stimulus induced changes in the raw voltage signal are apparent in a continuous recording from the experiment. **B)** Group average spectrograms are shown for each grating contrast level, color maps reflect percentage change in amplitude relative to the pre-stimulus period (black line indicates stimulus presentation). **C)** Group average normalized amplitude spectra (percent change) for each contrast condition averaged from the 250-500 ms post-stimulus time window (filled circles indicate peak amplitude frequency). Peak frequency of induced NBG for each subject across grating contrast levels (see Methods). Together B, C and D, clearly show a systematic increase in the NBG peak frequency with increasing contrast levels. **E)** Group mean amplitude time coarse for NBG and BBG ranges. While NBG shows a more sustained temporal profile, BBG is more typified by transient increases and stimulus onset and offset.

### NBG oscillation peak frequency is related to visual grating contrast

Previous work has suggested that changes in grating properties, such as contrast, influence the amplitude and frequency of NBG oscillations (18, 21, 25). As shown in Figure 2B, group averaged time-frequency plots clearly show that increases in grating contrast induce both an increase in NBG amplitude and peak frequency. These time frequency plots include both presented orientations, as they did not produce differing effects on NBG amplitude or frequency (i.e. no clear orientation selectivity for vertical or horizontal gratings was observed and therefore orientation is collapsed for all analyses below, see Figure S2). To more accurately evaluate the influence of grating contrast on NBG activity, we averaged values in the time window between 250-500 ms post stimulus onset (avoiding transient responses at stimulus onset and offset). Figure 2C shows the group mean normalized amplitude spectra for each contrast condition. With increasing levels of contrast, the amplitude and peak frequency of NBG oscillations increased. To obtain a more precise quantification of this effect and its variability, we identified NBG peak frequency at the single trial level and then averaged across trials (separately for each contrast level) and across electrodes (separately for each participant). This was performed to avoid any bias related to the different number of electrodes across individuals (see SI for additional control analyses). Specifically, grating contrast strongly influenced the NBG peak frequency, on average inducing a 4Hz shift in the peak frequency for each contrast level increment (mean peak frequency and standard error for 20% contrast: 36.4±1.4Hz; 50% contrast: 40.2±1.9Hz; 100% contrast: 43.8±2.03Hz; main contrast effects F(2,12) = 14.62, p=0.0006; Figure 2D & S3). Post-hoc comparisons confirmed that NBG peak frequency was statistically different between each contrast level (all p<0.05 Bonferroni corrected, from now p_corr_).

Peak amplitude was also significantly modulated by grating contrast, increasing on average from ∼250% to 400% signal change from low to high contrast (20% contrast: 249.7±54.5%; 50% contrast: 351.4±65.8%; 100% contrast: 405±66.2%; F(2,12)=22.3, p<0.0001, all post-hoc comparisons p_corr_ <0.05). Overall, these results confirm that the peak frequency in the NBG range varied as a function of contrast, increasing almost linearly within a 35-45 Hz range when subjects were viewing gratings with 20%, 50% and 100% contrast. However, it is important to note that this range reflects the average values, and individual subjects differed in their NBG frequency range (Figure 2D). Notwithstanding the variability in the peak frequency across individuals, the relationship between contrast and NBG peak frequency was detectable both at an individual and group level (Figure S3). In comparison, BBG, while increasing in amplitude across contrast levels (Figure 2B,C), did not display any characteristic spectral changes (see SI results).

### Gratings contrast can be decoded better using NBG than BBG

Given the strong influence of grating contrast level on induced peak frequency of NBG, we sought to test the reliability of this effect by decoding contrast levels using single trial normalized amplitude spectra over the NGB range for each electrode. Using a support vector machine with cross validation, and NBG spectra as the feature, grating contrast could be decoded significantly above chance for each subject (where chance = 0.33 and significance was assessed with permutation testing; see Methods). Overall, 76% of visually responsive electrodes showed significant classification accuracy (101/133 VEP sites). When using BBG as the feature, significant classification was achieved in only 35% of visually responsive electrodes (47/133). These data highlight a significantly greater number of electrodes that could successfully classify the stimulus contrast level when using NBG versus BBG (χ^2^(1)=44.41, p<0.001). Overall, the classification accuracy was higher for NGB signals versus BBG signals (considering all visually responsive electrodes: NBG median accuracy = 0.55, BBG = 0.39; Wilcoxon signed-rank test p<0.001; see Figure S4). To test the utility of our VEP selection of responsive sites, we also tested classification including non-VEP occipital electrode sites. These data displayed a similar trend, where 61% of electrodes (125/205; median accuracy 0.48) showed significant classification using NBG, and 25% of electrodes when using BBG (52/205; median accuracy 0.36). The anatomical distribution of classification accuracy closely matched the distribution of VEP sites shown in Figure 1 (Figure S4). More generally, electrodes outside of the occipital lobe did not show any significant classification (Figure S4). Together, these results highlight the stimulus dependence of visual NBG signals: grating stimulus attributes reliably shifted the NBG peak frequency, such that it was possible to reliably decode which grating was viewed based on single trial NBG spectra in early visual areas.

### NBG and BBG display different temporal responses to visual gratings

As is clear from Figure 2B, a notable difference between NBG and BBG was the temporal profile of response. Figure 2E shows the mean time course of NBG and BBG (after having averaged the normalized amplitude spectra within the 20-60 Hz and 70-150 Hz frequency ranges, respectively). NBG showed a sustained response throughout stimulus presentation, while the BBG response was transiently increased in amplitude at stimulus onset and offset. BBG did not return to baseline values between onset and offset, showing an additional sustained response, although of much smaller amplitude with respect to NBG (a temporal profile highly consistent to spiking activity under similar stimulus conditions (18)). In addition, Figure 2E suggests a slight delay in the onset of the sustained NBG, consistent with previous observations (19). To quantify these temporal dynamics, we calculated the relative onset time of NBG and BBG activity at the single trial level (see Methods). On average, NBG had an onset latency of ∼130 ms across conditions, while BBG onset occurred earlier, at ∼80 ms (NBG onset: 131.7±9.7 ms, BBG onset: 77.5±14.5 ms, t(6)=4.9, p<0.01). Intriguingly, the contrast manipulation had an opposite effect on the onset times in the NBG and BBG ranges. Higher contrast had a slightly later NBG response (20% contrast: 131.4±11 ms; 50% contrast: 128.1±11.4 ms; 100% contrast: 143.9±9.0 ms; F(2,12)=3.08, p=0.08; significant difference between 100% and 50% contrast levels, post-hoc comparisons p_corr_=0.029). Conversely, BBG onset latency was faster for higher contrast gratings (20% contrast: 99±14.1 ms; 50% contrast: 72.2±17.3 ms; 100% contrast: 67.5±14.6 ms; F(2,12)=10.83, p=0.002). In particular, the extremes of the contrast levels had significantly different BBG onset times (post hoc comparisons 20% vs. 100%: p_corr_=0.026). These opposite effects rule out the possibility that the onset time difference is simply related to a delay in detecting the onset when considering a slower frequency, as it would not explain why the stimulus attributes would influence the onset times differently in the BBG and NBG ranges.

In summary, NBG and BBG display differing temporal profiles of response to static grating stimuli (i.e., more sustained vs. more transient, respectively). Furthermore, they differed in their onset latency with the NBG occurring after BBG. This delayed NBG onset is far longer than the known latencies of visual cortex (e.g. as shown by the VEP N1 response above), but is consistent with previous reports (18, 19, 23). Having explored the spectral and temporal properties of gamma range amplitude, next we sought to quantify the phase consistency properties of induced gamma range activity across trials, within and between regions.

### NBG oscillations display variable phase clustering across trials

As noted above, and as shown in Figure 2, grating stimuli drive NBG oscillations of consistent frequency within individual’s, relative to the stimulus contrast. We therefore asked if these induced NBG oscillations are phase consistent across trials. Given that NBG onset latency occurred much later than the VEP, there may be a critical level of excitatory drive beyond which it is possible to observe NBG oscillations. In this sense, NBG might be related to a population resonance threshold, which may align oscillatory phases across trials. To assess if NBG oscillations displayed phase-locking during stimulus presentation, we computed a measure of phase consistency across trials (referred to throughout as inter-trial phase clustering, ITPC (26)). This measure is an index of the phase similarity across different trials at corresponding time and frequency points. ITPC was calculated for the different contrast levels separately. As shown in Figure 3A significant phase clustering was present at the onset and offset of the stimuli, in a broadband range spanning from low to high frequencies (2-160 Hz, occurring between 0 to 200 ms and 500 to 700 ms post-stimulus, p_corr_<0.05). Interestingly, no reliable ITPC was found throughout stimulus presentation in the NBG range, during which NBG amplitudes are maximal. Thus, across trial phase consistency was present only at stimulus onset/offset, while no consistent phase similarity across trials was maintained during stimulus presentation. Given the transient and broadband nature of the ITPC, VEP potentials at stimulus onset and offset are suspected to be a contributing influence. It is for these reasons, along with its non-oscillatory nature, that BBG is not amenable to targeted phase analysis. While NBG phase dynamics may temporally vary across trials, these oscillations may still be phase consistent with other brain regions within trials. Therefore, we next tested whether the phase relationship between electrodes would be consistent during stimulus presentation.

**Figure 3.**
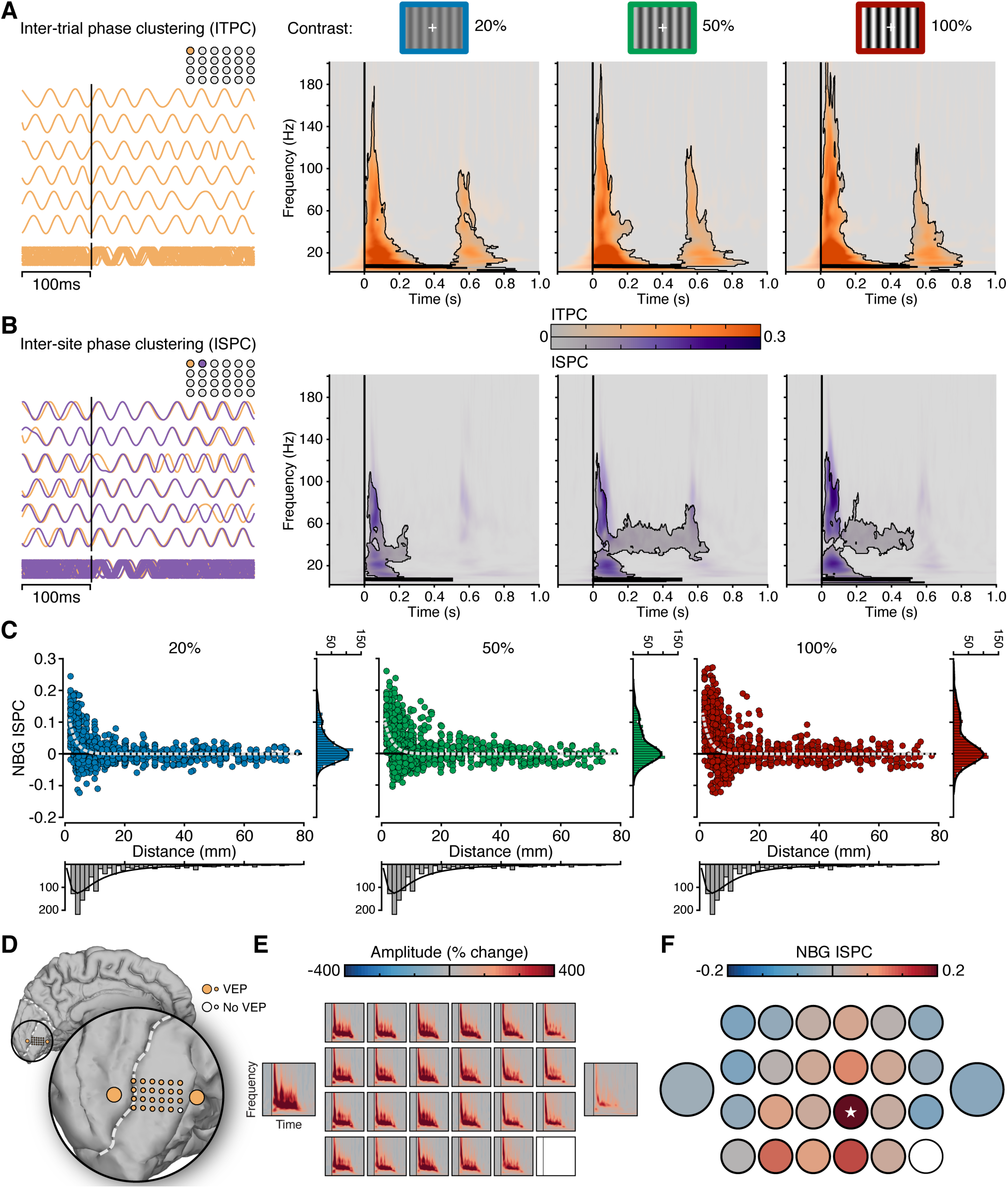
Inter-trial and inter-site phase properties of NBG. **A)** On the left, examples of NBG phase across repeated trials for a single electrode site, used for estimating inter-trial phase clustering (ITPC). On the right, time-frequency plots show group mean ITPC for each contrast condition. ITPC shows broadband clustering for stimulus onset and offset for each contrast level. **B)** Similar to A, but for two example electrode sites, used for estimating inter-site phase clustering (ISPC). Time-frequency plots show group mean ISPC (250-500 ms) for each contrast condition (based on all valid electrode pairs n = 1252, see Methods). ISPC shows broadband clustering for stimulus onset followed by sustained NBG ISPC predominately for the 50% and 100% contrast levels. ITPC and ISPC reflect normalized values relative to the pre-stimulus period. **C)** NBG ISPC is shown for each unique electrode pair as a function of inter-electrode distance for each contrast condition (see methods). Data are fitted with a single term exponential decay function, using nonlinear robust fitting (dashed line). **D)** Electrode location of macro- and mini-ECoG grid in subject N7. **E)** Spectrograms show the mean amplitude response across the electrodes shown in D, for the 100% contrast level. **F)** NBG ISPC (100% contrast) for the same electrodes in D,E, relative to a seed electrode site (indicated with a white star). Despite strong responses across electrodes, NBG ISPC is focal and has a rapid decay within the mini-grid.

### NGB oscillations display local phase clustering between sites

One of the central theories of the functional role played by gamma oscillations is the formation of a communication channel between regions by rhythmic synchronization or coherence (27). To test for evidence of phase consistency across visual cortex in the NBG range, we computed a measure of phase-based consistency (referred to throughout as inter-site phase clustering, ISPC (28)) across all pairs of electrodes within each subject. This measure is an index of the stability in the relative phase relationship between two cortical sites (phase difference) across different trials at corresponding time and frequency points. As shown in Figure 3B, ISPC was present at stimulus onset, in a broadband range spanning from low to high frequencies, similar to the ITPC pattern (5-120 Hz, occurring between 0-200 ms, p_corr_<0.05). In addition, there was evidence of ISPC in the NBG range, sustained throughout stimulus duration. This pattern of NBG ISPC was weaker than the stimulus onset response, but significantly higher than baseline when considering the higher contrast conditions (between 30-60 Hz and occurring from ∼100 to 500-600 ms post-stimulus).

Importantly, inter-regional coherence might be strongly dependent on the distance between recordings sites, thus difficult to detect at larger cortical distances or when averaging across different scales (mm vs cm), as done in Figure 3B. We next exploited the high-density of the electrode arrays used in this study to investigate the relation between phase consistency and inter-electrode distances. Figure 3C shows NBG ISPC for all electrode pairs (n = 1252) as a function of electrode pair distance for the three contrast levels (NBG ISPC was averaged within a 250-500 ms time window and the 20-60 Hz frequency range, consistent with all analyses above). To quantify this relation, we fitted an exponential decay function to the NBG ISPC over distance and obtained similar decay dynamics for the three contrast levels, with a spatial decay constant (distance needed to observe a 1/*e* drop of phase-consistency values, i.e. a 37% decrease) ranging from around 2.3 to 2.5 mm (Figure 3C; see SI and Figure S5 for a comparison of the decay rate computed using other phase-based synchrony measures). Consistent with previous work in non-human primates, these data suggest that induced NBG is synchronous locally, for distances less than 5 mm (19). To confirm that the rate of decay observed was reliably present also when considering single-subject data, we report an example of a mini-ECoG grid array in subject N7. The mini-ECoG array displayed clear NBG responses to visual gratings (Figure 3D and E) while phase consistency was present only at neighboring electrodes (Figure 3F) within 2-6 mm. These results support a very local effect of phase-based coherence during the viewing of gratings: although many cortical locations exhibit NBG responses to gratings, only those in the range of few millimeters can reliably display a consistent phase relationship.

### Summary of NBG and BBG responses to grating stimuli

Altogether, the results from the visual grating task suggest that grating stimuli induce robust NBG oscillations. However, replicating previous work, increasing contrast levels of grating stimuli increased the frequency of NBG oscillations. This effect of contrast on NGB peak frequency is consistent across individuals, although the specific frequency range differs, as often the case when examining inter-individual variability of cortical rhythms (29). NBG displayed a sustained time course during stimulus presentation, with a delayed onset of ∼130 ms. This response is in contrast to BBG activity which showed a more transient response at stimulus onset and offset, with a shorter onset latency of ∼80 ms. This temporal profile of BBG is highly consistent with spiking activity recorded under similar conditions (18). The sensitivity of NBG oscillations to the properties of grating stimuli has recently been interpreted as a dependence – whereby other classes of visual stimuli do not reliably induce NBG oscillations. Furthermore, NBG displayed no consistent phase alignment across trials. However, NBG did show phase consistency between nearby recording sites, which exponentially decayed with inter-electrode distance. Beyond these observations, a central point of a recent debate (30, 31) has focused on whether these types of NBG responses occur for more complex naturalistic stimuli. We therefore next quantified the gamma range responses in visual cortex of the same subjects during the viewing of natural image stimuli.

### Natural image stimuli induce BBG but not reliable NBG responses

It is currently debated whether NBG oscillations are consistently induced during the viewing of natural images, with evidence supporting both possibilities (30-33). To test for the presence of NBG in response to natural images, we presented grayscale images of different visual categories (faces, houses, bodies, limbs, cars, words, numbers, phase-scrambled noise) to a subset of our subjects (5 out of 7). Images were presented for 1 sec and subjects were asked to perform an exemplar 1-back task responding by a button press to stimuli repeated back to back. Again, raw voltage responses suggested clear stimulus induced activity in visual cortex, but with more varied signatures, as shown in Figure 4A. We next quantified the time-frequency representation of these responses across subjects. Figure 4A also shows the group mean spectrograms for each stimulus category. Time-frequency maps show the clear presence of a BBG response at stimulus onset, spanning from ∼20-200 Hz. This feature slowly weakened after stimulus onset, but showed a minor increase again at stimulus offset. Strikingly, there is a lack of sustained NBG response to any of the visual categories presented with comparable spectral characteristics to experiment 1. Otherwise, the responses recorded in early visual cortex were highly stereotyped across the different image categories, despite the diversity of image content. These observations were consistent for both macro- and mini-ECoG electrodes (Figure S6). It is also important to note that these images were matched for several low-level attributes (see Methods (34)).

**Figure 4.**
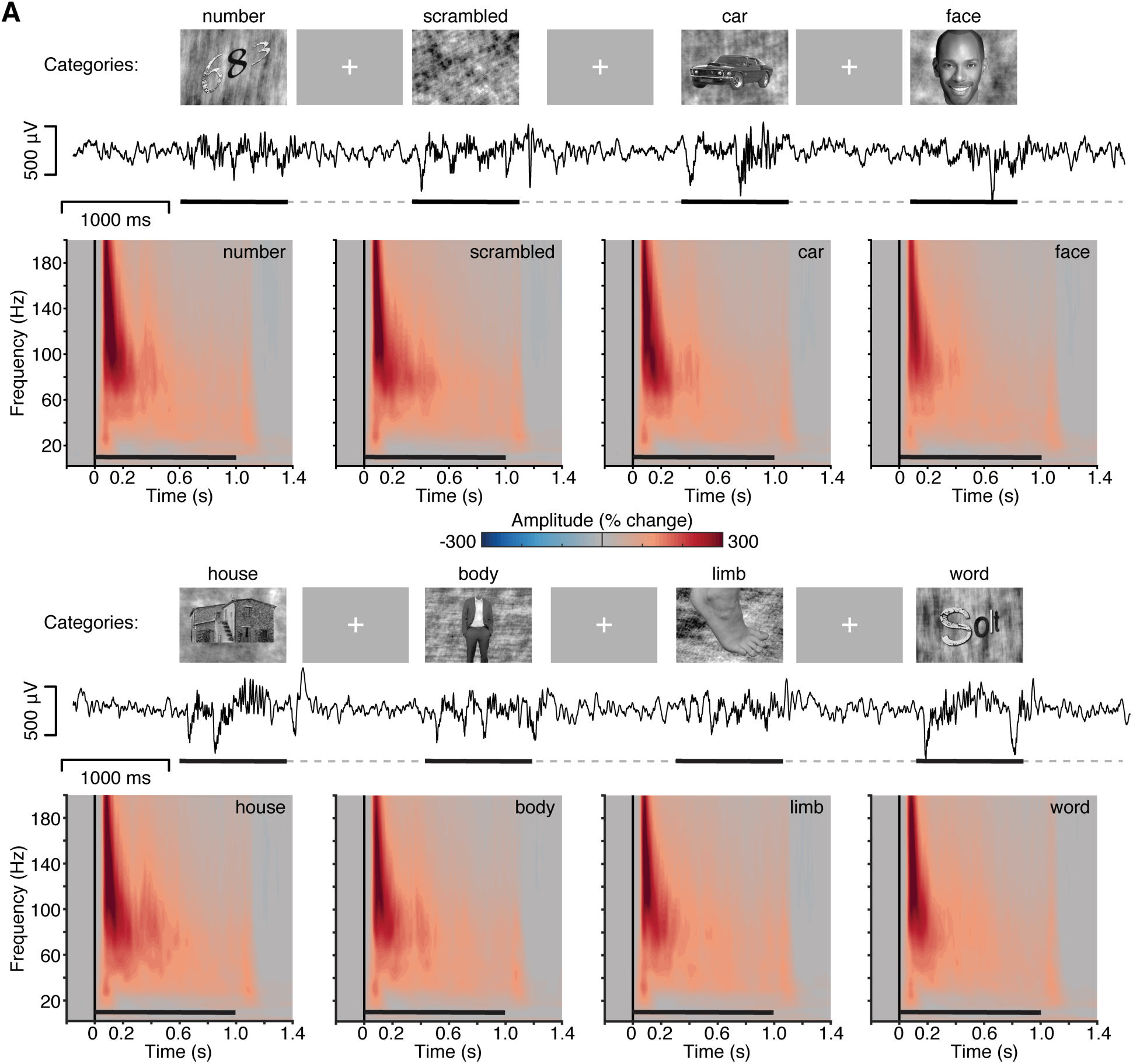
Experiment 2, spectral response to natural image stimuli. **A)** Experiment 2 stimuli, example voltage response and group mean spectrograms. Stimuli were gray scale images from eight visual categories (number, scramble, car, face, house, body, limb and word), presented for 1000 ms (black line), with a random inter-stimulus interval (ISI; dashed line) of 1-1.5 s (see Methods). Example voltage response to stimuli is shown (subject N6, same as Figure 1A). Mean group spectrograms show a highly consistent time-frequency response profile, typified by a strong BBG response to stimulus onset (color maps reflect percentage change in amplitude relative to the pre-stimulus period; black line indicates stimulus presentation).

Next, we computed the power spectra to obtain a clearer comparison between the spectral composition of the signals in visual cortex in response to natural images *vs.* gratings (Figure 5). By directly contrasting the two task responses in the same sample and using the same analysis parameters, our results showed a different spectral pattern in early visual cortex for the viewing of natural images compared with gratings. As shown in Figure 5A, BBG responses were clearly visible in the power spectra for natural images, while no peaks in the NBG range were observed. The mean spectral response to different categories were approximately indistinguishable from one another. Conversely, NBG peaks were clearly visible on top of a broadband power increase for grating stimuli (Figure 5A). Differences in normalized amplitude spectra may be due to different baseline conditions. However, similar spectral differences were also apparent when considering the non-normalized power spectra (Figure 5B), which also shows that the baseline recordings (i.e., when no visual stimulus was on screen) were similar across experiments.

**Figure 5.**
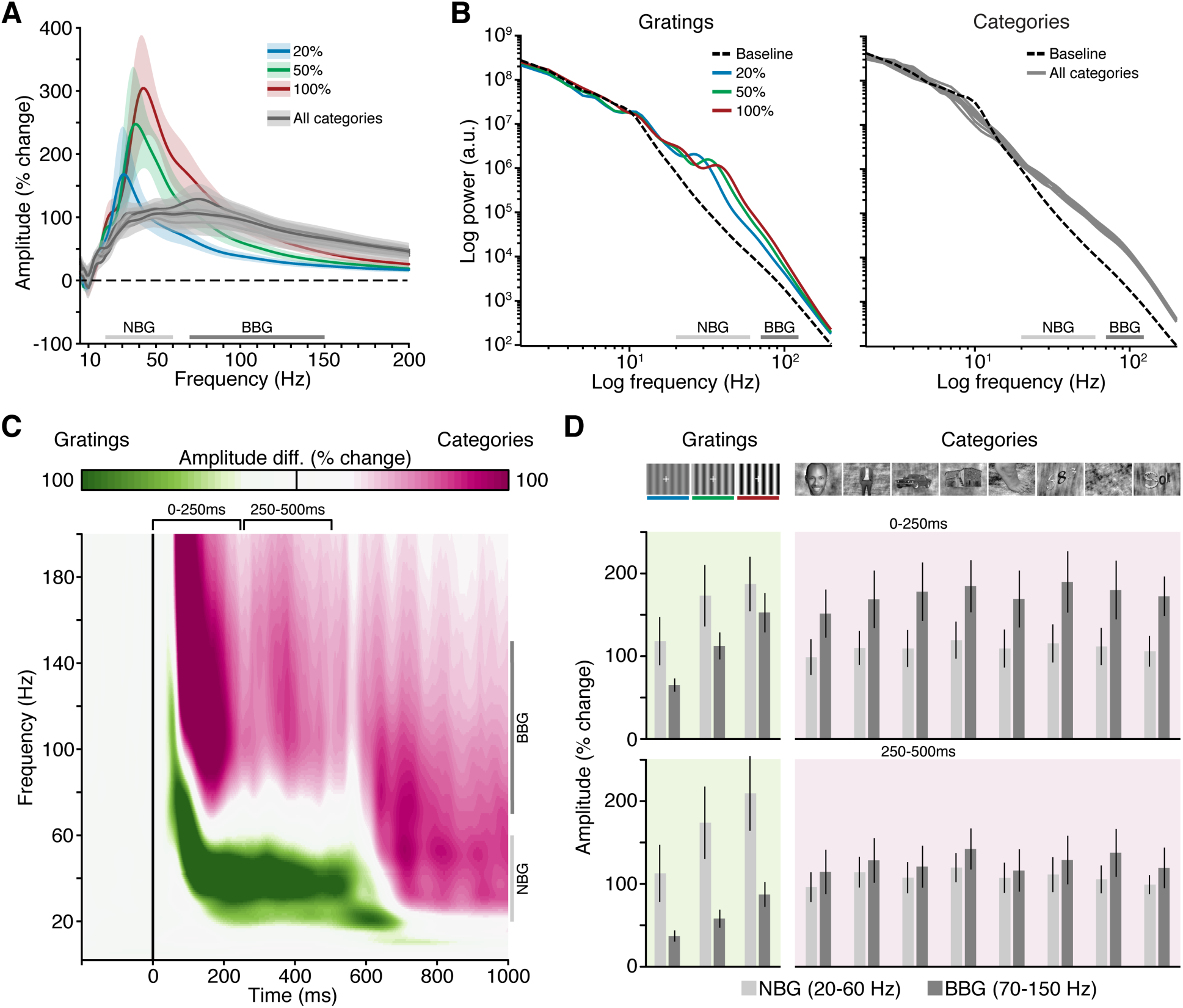
Experiment 1 & 2 spectral comparison. **A)** Mean group normalized amplitude spectra (% change) are show for all conditions across both experiments (image categories are given same grey color given highly overlapping data), shading reflects standard error of mean. **B)** Mean group power spectra (log-log axis) are shown for both experiments, including mean baseline power spectra. **C)** Mean group ‘task selectivity’ spectrogram (grating *vs*. categories). Time-frequency maps highlight magnitude of response relative to task (green for time-frequency points larger for gratings; purple for time-frequency points larger for categories). Conditions in both tasks have been collapsed (note: gratings are presented for 500 ms and natural images for 1000 ms). **D)** Mean group amplitude response for NBG and BBG for early (0-250 ms) and late (250-500 ms) time windows as indicted in (C). Across A, B and C, NBG and BBG range is indicated for reference. Together these plots highlight that BBG changes extend through the classical NBG range, therefore amplitude increases in the NBG range cannot be immediately inferred to be increased NBG oscillatory power.

To more precisely capture these task differences in the time-frequency domain, we constructed a ‘task selectivity’ spectrogram as shown in Figure 5C. For each recording site, we computed the difference in mean amplitude (% change) values between gratings (experiment 1; all conditions) and natural images (experiment 2; all conditions). We then depicted the mean time-frequency task differences across all recording sites to highlight the time-frequency features that were more strongly driven by each experimental stimulus. As shown in Figure 5C, the task selectivity spectrogram recapitulated the stimulus class-dependence described above for NBG and BBG, with gratings driving stronger NBG range responses, and natural images driving stronger BBG responses. However, while this comparison helps to emphasize task differences, it is important to note that both experiments drove amplitude increases in NBG and BBG frequency ranges, but such increases reflect highly different response signatures as clearly observed in Figure 5A and B. Indeed, it is clear that the response to natural images is not confined to the frequency range >70 Hz, but actually spans a broader range falling as low as 20 Hz. This becomes particularly evident considering the time points where only categories are on screen (Figure 5C; note that gratings were presented for 500 ms, natural images for 1000ms). Figure 5D highlights this point by showing the group mean amplitude increases for NBG and BBG across all experimental conditions for early (0-250 ms) and late (250-500 ms) time windows. As is evident in Figure 5D, NBG and BBG are elevated above baseline for all stimuli at early and late time windows. Critically, increases in NBG for grating stimuli are reflective of a spectral peak, whereas for natural images it is simply part of a broadband amplitude increase extending into the NBG range.

For these reasons, caution must be taken when simply quantifying amplitude or power changes within this NBG range as synonymous with shifts in an oscillatory signal, as recently argued by Haller et al. (35). To deal with these issues of spectral overlap, and genuine changes in NBG oscillations, we employed a further fitting procedure to automatically identify oscillatory peaks above the broadband aperiodic component in the power spectrum (35), similar to prior studies of NBG (22). NBG oscillations, while prominent for the visual gratings, were not present (or not detectable) for natural images, as it was not possible to model the responses as containing one or more oscillations in the 20-150 Hz range (see SI). This recapitulated our observations that grating stimuli produced a readily identifiable NBG peak, while no oscillatory-like response could be detected for responses to the natural image categories presented. In summary, using a wide variety of visual categories we found no reliable evidence of NBG responses of comparable power and frequency content as those recorded in response to visual gratings, supporting a stimulus class-dependent dissociation of the spectral characteristics of early visual responses.

## Discussion

Using high-density intracranial recordings from early visual cortex in the human brain, we observed distinct response properties of NBG and BBG to grating and natural image stimuli. NBG and BBG reflect the historically defined gamma and high-gamma ranges, respectively.

For grating stimuli, NBG showed a sustained response throughout the stimulus duration, with a delayed onset latency (∼130 ms). BBG showed a transient response to stimulus onset and offset, with a more rapid onset latency (∼80 ms). Across increasing levels of grating contrast (20%, 50%, 100%), the peak frequency of NBG increased (∼36 Hz, 40 Hz, 44 Hz). BBG showed no reliable changes in frequency characteristics. Both NBG and BBG showed increasing response amplitudes with increasing contrast levels. Together, these response features supported successful classification of stimulus contrast, where classification accuracy was significantly greater for NBG compared to BBG. Across repetitions of grating trials, NBG displayed no consistent phase alignment. However, NBG did show phase consistency between nearby recording sites, which rapidly decayed with inter-electrode distance. While broadband transients were seen for inter-trial and inter-site analysis, meaningful interpretation of BBG phase properties is problematic and potentially biased by event related potentials.

For natural images, no reliable NBG responses were observed across the eight image categories examined. In contrast, robust BBG responses were observed for all image categories, being transiently maximal at stimulus onset. Together, these findings clearly dissociate NBG and BBG activity in spectral, temporal, and functional domains. These differences have implications for functional mechanisms ascribed to NBG and BBG, and more broadly to experimental design and data analysis, as discussed below.

### Narrowband gamma activity

NBG oscillations have been proposed as a mechanism for coordinating local and distal spiking activity and support communication in neocortex (2). Consistent with this view, extant evidence suggests NBG in early visual cortex influences local spiking activity (36), can synchronize with downstream targets in the visual system (37, 38) and is modulated by attention (6, 38). Criticisms of this mechanistic view have focused on challenging the ubiquity and reliability of NBG oscillations (39). If NBG is an important mechanism for visual perception, it should be a robust response feature that is ubiquitous across visual stimulus inputs. However, a growing literature has emphasized the strong stimulus dependence of NBG in two ways. Firstly, NBG displays an attribute dependence, where properties of NBG, such as amplitude and frequency, are dependent on properties of the driving stimulus. This is exemplified by the data we report on the relationship between NBG peak frequency and grating contrast level (Figure 2). This grating-contrast dependence was highlighted by Ray and Maunsell (18) in non-human primate V1, but has also been reported elsewhere in human V2/3 (21), and can be detected using non-invasive methods (e.g. 25).

An additional line of criticism has focused on the stability and reliability of induced NBG. These studies have highlighted how NBG lacks reliable signal properties, such as phase stability, to support neural communication at the time scales associated with perception and cognition. For example, NBG may lack the ‘clock’ like attributes to synchronize distributed neural circuits (40-42). Importantly, these criticisms have been challenged, whereby NBG oscillations may still fulfill their functional role with variable phase and frequency attributes (43, 44). Our results show some local phase consistency in the NBG range, although restricted to adjacent early visual locations within a few millimeters from each other.

Secondly, NBG displays a class or category dependence, such that NBG responses are more reliably observed for specific types of visual stimuli. For example, NBG is highly responsive to grating like stimuli (e.g. Gabor patches, checkerboard stimuli), but is often reduced or absent for more complex stimuli such as noise patterns and natural images (39). Earlier work in cat V1 highlighted this dependence (45), which more recently has been detailed and extended in human primary visual cortex by Hermes et al. (22). However, the principles governing this NBG stimulus dependence are subject to debate (30, 31) as some complex images may produce NBG responses (22, 33). Adjudicating these findings, specifically for natural images, is further compounded by NBG sensitivity to color (46). These findings, together with our own observations, suggest that, in some cases, NBG responses to complex stimuli could be explained by their sensitivity to low-level stimulus attributes intersecting with the receptive field.

While our observations are consistent with previous work suggesting NBG is sensitive to stimulus properties, there are several criticisms to consider. Firstly, our measurements may not have been sensitive enough to capture NBG responses to natural images. This explanation seems unlikely as our electrodes (macro-/ mini-ECoG) are of similar or smaller dimension to previous ECoG studies in the macaque (33), and show responses highly similar to intracortical recordings under similar stimulus conditions (18). Secondly, we did not enforce fixation during the second experiment, thus eye movements could have played a role in the different responses between experiments. However, previous work has reported NBG oscillations during free viewing of natural images (33), limiting eye movement as a critical confound. A third criticism is related to the use of average responses (across individual stimuli within image categories): as stimuli were only presented once, this may have potentially blurred putative NBG oscillations with different peak frequencies. While a possible confound, previously observed NBG responses to natural stimuli did not show a broad variability of peak frequency that would be lost in averaging data (33). This is further supported by the uniformity of mean time-frequency responses we observed across categories despite the wide variability of stimuli. Finally, in our experiment, natural image stimuli were grayscale whereas prior work, reporting NBG responses, used color images, suggesting NBG may be sensitive to chromatic stimulus features. Indeed, recent work has shown NBG is enhanced particularly by red and orange hues, even for uniform color stimuli without any grating like or patterned structure (46). In this respect, color may present an additional domain of NBG stimulus dependence rather than invariance.

### Broadband gamma activity

BBG activity takes many names in the literature such as ‘high-gamma’, ‘high-frequency activity’, ‘high-frequency broadband’ or ‘broadband’ (13). Research utilizing human intracranial recordings has particularly focused on this signal given its many desirable properties as a marker of local neocortical response (14). Spectrally, BBG is commonly observed as a wideband non-oscillatory signal. As shown in this study and many others, the time course of BBG is highly similar to population spiking or multi-unit activity under similar task conditions or simultaneous recording (15-17). Indeed, BBG responded transiently to visual stimuli in a manner consistent with MUA. Despite the strong apparent temporal correlation between BBG and MUA, further work is required to elucidate the biophysical generators of this response. Clarifying these relationships may allow for improved neurophysiology interpretation of BBG and related responses, including better insight into neural population dynamics. For example, recent evidence from Leszczynski et al. (47), suggest some dissociation of BBG and MUA when considering their laminar distribution. Importantly, one practical implication of these BBG signal properties and potential generators is the uncertainty in applying ‘synchrony’ type measures to this frequency range, or interpreting power changes in the BBG as changes in an oscillatory signal. Overall, while a more detailed understanding is required, the BBG signal serves as reliable marker of electrocortical response closely associated with population activities proximal to the electrode site (48).

While our definition of BBG is consistent with previous studies of high-gamma activity (70-150 Hz), such ranges are often variable in the literature (13). It is important to note that defining the band pass of BBG as 70-150 Hz is more a practical convenience than a formal definition. As Miller et al. (14, 49) have argued, BBG is likely a wide band phenomenon that can extend into lower frequencies, overlapping with the NBG range. This overlap is clearly seen in our recordings as shown in Figure 5, and as noted above, can confound analysis and interpretation of gamma amplitude/power increases as oscillatory responses. For these reasons investigators have appropriately used more data driven methods to model or decouple oscillatory and broadband spectral responses (22, 35, 49, 50). Improving data driven and physiologically informed decomposition of frequency domain analysis is therefore a critical area for development in cognitive and systems neuroscience.

### Implications for theory and experiment

As indicated above, the stimulus dependence of NBG has been invoked as a challenge to its utility as a mechanism of visual perception and as a vehicle of cortical communication more generally (22, 31, 39). While compelling theoretical and experimental arguments have been advanced to deal with some of these issues (e.g. (43, 51)), future computational models will need to address the growing diversity of parametric relationships between visual stimuli and NBG response properties.

Beyond these theoretical issues, what are the more applied implications of these findings? Our work highlights that NBG and BBG have distinct spectral, temporal and functional properties, which together may help to reinterpret prior work. An intriguing example relates to past efforts to elucidate the neural correlates of blood oxygen level dependent functional magnetic resonance imaging (BOLD fMRI). In a landmark paper, Logothetis et al. (52) compared electrophysiological responses (spiking, MUA, LFP) to BOLD fMRI using a sophisticated experimental system allowing simultaneous acquisition of these data. They found the LFP signal showed the best temporal correlation with the BOLD response, being sustained during stimulus presentation of different durations unlike the more transient local MUA or single unit spiking activity. In simple terms, this finding was interpreted as indicating that the BOLD signal reflects the consequence of local input to a given brain region, and not its spiking output (52). However, these measurements were obtained from early visual cortex, using checkboard stimuli, which drove robust NBG responses. As observed here, these NBG responses were sustained throughout stimulus presentation of different durations. Critically, the LFP was defined as power in the 40-130 Hz frequency range, which captured both NBG and BBG activity (52). It is therefore of great interest to consider how the stimulus dependence of NBG, its sustained response properties, together with the partial correlation of BBG and MUA, could provide a new interpretation of these pioneering data. In particular, the strong LFP-BOLD correlation may be less robust across other stimulus classes in visual cortex, and the LFP range employed may not be easily dissociated from MUA under these conditions. This interpretation is supported by the strong correlation observed between LFP/MUA amplitude modulations and BOLD activations, and related human intracranial studies inferring a close relationship between BBG, spiking activity and BOLD fMRI (17, 53). This link between BOLD fMRI and BBG has received several replications across multiple brain networks (54). Given the debate surrounding these relationships (55), the interpretation offered here may in part reconcile prior discrepancies.

### Conclusion

Our findings build upon and integrate prior work in the human and non-human primate brain (18, 22), and suggest that a more careful theoretical and empirical approach to gamma range activity is required. Spectral, temporal, and functional dissociation of narrowband and broadband gamma activities suggest critical differences in these signals, and warn against analyses/interpretations that may conflate the two. Of particularly note, the stimulus dependencies of NBG, if further supported by future work, suggests a more restricted and nuanced functional role of neocortical gamma oscillations in visual cortex. Generalizing these findings to gamma activity in other cortical / non-cortical structures (e.g. hippocampus), or species, should be done cautiously as a diversity of brain circuits generate behaviorally relevant gamma range responses (56, 57). Indeed, while future work is required to adjudicate these more consequential mechanistic inferences, the robust stimulus-response properties of NBG may still provide a powerful methodology for studying visual circuit properties. In this sense, neocortical NBG oscillations may help us understand the mechanisms of vision, without being a fundamental mechanism for vision.

## Materials and Methods

### Subjects

Intracranial recordings (electrocorticography, ECoG) were obtained from 7 subjects (N1-7; 6 males, mean age 36 years, ranging from 20-54 years) undergoing invasive monitoring for the potential surgical treatment of refractory epilepsy at Baylor St. Luke’s Medical Center (Houston, Texas, USA). Subject information is detailed in Table S1. All subjects provided written and verbal voluntary consent to participate in the experiments reported here. All experimental protocols were approved by the Institution Review Board at Baylor College of Medicine (IRB protocol number H-18112).

### Electrode Arrays

All reported data were recorded using subdural cortical surface electrode strip arrays (PMT corp., MN, USA). Electrode arrays were custom designed to incorporate mini-ECoG electrodes (0.5 mm diameter) into a standard macro-ECoG (2 or 3 mm diameter) clinical array configuration (58). The standard clinical ECoG strip array has 8 macro electrodes linearly arranged with a center-to-center distance of 10 mm. The mini/macro hybrid arrays used in this study had two different configurations: for hybrid array A, four mini electrodes where positioned around the first 4 macro electrodes; for hybrid array B, a high-density grid (4 x 6) of mini electrodes was positioned between the first two macro electrodes (in this configuration the distance between the first two macro electrodes was modified to be 18 mm). The two arrays are represented in Figure 1A. For subject electrode information see Table S1 and Figure S1.

### Electrode Localization

Electrode locations were determined by first co-registering a post-operative CT scan to a pre-operative T1 anatomical MRI scan for each subject, using FSL and AFNI (59, 60). The cortical surface location of each electrode was based on the local projection of electrode coordinates (identified as clear hyperintensities on the aligned CT) to a cortical surface model reconstructed from the anatomical T1 scan using Freesurfer (version 5.3; (61)). Mini-ECoG electrode coordinates were often not clearly identifiable from the CT image and their position was therefore calculated with custom functions combining the macro-ECoG coordinates with the known array geometry. Electrode locations were projected onto each individual cortical surface model using AFNI/SUMA and visualized using iELVis software functions (62) in Matlab (v2016a, MathWorks, MA, USA). For the current study, electrodes of interest where those localized to the occipital lobe.

Anatomically, the parieto-occipital sulcus served as a dorsal boundary and the lingual gyrus as a ventral boundary. Laterally, the trans-occipital sulcus and posterior aspect of the superior temporal sulcus served as dorso-lateral boundaries. Anatomical identification of occipital sites was performed with respect to each individual electrode localization. As detailed below, occipital electrodes were further sub-selected by means of a functional response criterion (Figure 1B). To visualize all occipital electrode locations in a common space, each electrode coordinate was transformed into the Talairach coordinate system and represented on the Colin N27 brain (Figure 1C). To do so, each subject’s T1 scan was transformed to match the template using AFNI, and the same transformation was applied to the electrode coordinates.

### Experimental tasks

All experiments were performed at the bedside in a quiet and dimmed patient room. Subjects performed two visual tasks (with the exceptions of subjects N1 and N2, who only completed the visual grating task, described below). For both tasks, stimuli were presented on an adjustable monitor (1920×1080 resolution, 47.5×26.7 cm screen size, connected to an iMac running OSX 10.9.4) at a viewing distance of 57 cm (such that 1cm = ∼1 deg visual angle). Both tasks were programed using Psychtoolbox functions (v3.0.12) (63) running on Matlab (v2017a, MathWorks, MA, USA).

### Visual grating task

In the visual grating task, participants (all 7 subjects) were shown full screen static grayscale grating stimuli – subtending ∼25° of visual field. Grating stimuli had a sine wave spatial frequency of 1 cycle/degree to maximize V1/2 responses (64) and were presented for 500 ms with a random inter-stimulus interval (ISI) between 1500-2000 ms (see Figure 2). The timing of the stimulus presentation was selected in order to limit the influence of afterimage effects. The contrast and orientation of the stimuli were manipulated experimentally in the following way: 3 levels of Michelson contrast (20%, 50% & 100%), and two orientations (0° & 90°). There were 30 trials for each contrast level, with equal numbers of vertical/horizontal orientation. The participants were required to maintain fixation (marked by a cross) and respond (via button press) whenever a target oddball stimulus was presented, which was a randomly occurring grating stimulus with a 45° orientation (15 targets in total, 5 for each contrast level). Performance was monitored by an experimenter present in the patient room. These target trials were not included in data analysis. Overall, the total number of trials presented was 105 with the task lasting around 6 minutes. The contrast levels were selected to maximize the perceptual and neural separation of stimuli and putative gamma effects, respectively, and to be similar to previous work (18). Based on previous work and our own pilot studies, this range is optimal given that gratings below 20% contrast often fail to reliably produce gamma band responses (18, 21). As grating contrast was the main manipulation of interest, all data analyses were collapsed across the two orientations (see Figure S2).

### Visual category task

In the visual category task, subjects (subjects N3-7) were presented grayscale images from 8 visual categories (faces, houses, bodies, limbs, cars, words, numbers, phase-scrambled noise) in random order (varying in position and size within a bounding box of phase-scrambled noise, subtending 15° degrees of visual field; see Figure 4). Visual stimuli were selected from a publicly available corpus that has been successfully used as a visual category localizer in human fMRI studies (34). Stimuli were presented for 1000 ms, with a random inter-stimulus interval between 1000-1500 ms. Subjects were required to respond (via button press) whenever they detected a specific stimulus being repeated back to back (1-back task). As for the first experiment, performance was monitored by an experimenter present in the patient room. The target trials were discarded from the analysis. 15 different stimuli were presented for each category and 10 random images were repeated (serving as targets), leading to a total of 130 trials. On average the task was 7 minutes in duration.

### Electrophysiological Recording

ECoG signals were recorded at 2kHz, with a bandpass of 0.3-500Hz using a 128 channel BlackRock Cerebus system (BlackRock Microsystems, UT, USA). Recordings were referenced to an inverted subdural intracranial electrode (i.e., touching the dura). Stimulus presentation was continuously logged via a photodiode sensor (attached to monitor) synchronously recorded at 30kHz. Photodiode recordings were also performed to check for any subthreshold stimulus flicker (see SI).

### Preprocessing

All signal processing was performed using custom scripts in Matlab (v2016a). Raw ECoG signals were imported into Matlab and visually inspected for the presence of line noise, recording artifacts and interictal epileptic spikes. Any epileptic or artefactual channel was excluded from further analysis. Each channel was notch filtered (60 Hz and harmonics) and re-referenced to the common average of all artifact-free macro-ECoG electrodes. Mini-ECoG electrodes were not included in constructing the common reference signal given that their spatial density could potentially bias the reference signal toward visual responses. Next, we employed a functional criterion to identify responsive electrodes within early visual cortex based on the presence of a visual-evoked potential (VEP). VEPs were constructed by averaging all trials (pre-processed data) for the grating task. The resulting VEP was defined as ‘responsive’ if it passed two criteria: 1) the standard deviation for the response window (0-250 ms, to capture the common N1/P1 response) was at least three times greater than the standard deviation for the baseline window (−1000-0 ms); 2) the voltage range for the response window was at least 10 times larger than the voltage range for baseline window. Overall, 65% of the anatomically identified occipital cortex electrodes survived this functional inclusion criterion (133 out of 205 total electrodes, ranging from 36-97% at the individual level) see Figure 1 and Table S1. This functional criterion was employed to exclude electrodes that were not corrupted by noise but still lacked a robust signal (e.g. not making good contact on the cortex, or outside early visual areas despite anatomical selection). We used the VEP in order to limit selection bias with respect to the presence/absence of the signal of interest (i.e. the presence of narrowband or broadband gamma).

### Spectral analysis

Signals from selected electrodes (i.e., continuous voltage time-series filtered and re-referenced) were down sampled to 1kHz and convolved with a family of Morlet wavelets, with central frequencies ranging linearly from 2 to 200 Hz in 1 Hz steps (7 cycles). The magnitude and angle of the complex convolution result were used as instantaneous amplitude and phase estimates as detailed below. Based on a large literature from human and non-human primate studies, narrowband gamma (NBG) was defined as neural activity between 20-60 Hz and broadband gamma (BBG) as neural activity between 70-150 Hz. The instantaneous amplitude was normalized into a percent change signal by applying a baseline correction at each time-frequency point (baseline division, using the average pre-stimulus values −500ms to 0ms for each trial). We first explored the time-frequency percent change maps averaged across trials separately for each task-specific condition (visual grating task: contrast levels, time window −200 to 1000 ms around stimulus onset; visual category task: category types, time window −200 to 1400 ms. For both tasks, two main measures were derived from the time-frequency percent change values: the normalized amplitude spectra (obtained by averaging across the time dimension, using the 250-500 ms post-stimulus window – to avoid the transient response to stimulus onset; see Figure 2C,D) and the time course of NBG and BBG (obtained by averaging across the frequency dimension, using the 20-60 Hz and the 70-150 Hz ranges respectively). Additionally, power spectra for each task condition were also computed (squaring the non-normalized amplitude values for the single trial data and then averaging across time using the same windows described for the normalized amplitude spectra above).

#### NBG Peak frequency

We first tested whether the frequency of NBG was modulated by the contrast level of visual grating stimuli (20%, 50% & 100%). The single-trial normalized amplitude spectra were used to identify the frequency displaying the highest amplitude change in the narrowband gamma range (NBG, 20-60 Hz). Trials lacking a clear frequency peak were discarded (∼2% of trials). The identified peak frequency and amplitude values were averaged across trials (separately for each contrast level) and across electrodes (separately for each participant). The average values (for both peak frequency and amplitude) were analyzed by means of a within-subject ANOVA, modelling the contrast levels as predictor (modeled as fixed effect, subjects as random effects) using R statistical software (v3.4.3; (65)). We performed two additional control analysis: we computed the peak frequency on the average normalized amplitude spectra (on the electrode average vs. the single trials) and repeated the above ANOVA, to control that the single-trial were able to capture the peak frequency values as reliably as an average signal with better signal to noise ratio. We also performed a mixed-effects analysis, in which we did not average across electrodes, but we modeled them as random effects. Both analyses are reported in the SI and they are in agreement with the main analysis reported in the result section.

#### NBG and BBG onset latency

Next, we evaluated the onset latency of NGB oscillations. We computed the onset time of the single trial NBG amplitude change by adapting a previously published method to our data (54). First, for each NBG signal, we marked the first time point, at which the NBG amplitude exceeded a threshold (> 75th percentile for at least 40 ms in the time window between −100 to 500 ms; 15% of the trials were discarded as they did not meet this criterion). After that, a 300 ms wide window was built around that time point (200 ms before and 100 after). This window was segmented into 50 ms bins with 20% overlap and a linear regression was fit to each bin. The first time-point of the bin with the highest slope and smallest residual error was defined as the onset of the NBG. The onset time values were averaged across trials and across electrodes and were analyzed by means of a within-subject ANOVA (as described above for the peak frequency analysis). The same procedure was repeated to estimate BBG onset latency.

#### NBG phase synchrony across trials and locations

To assess if NBG oscillations displayed phase-locking during stimulus presentation, the instantaneous phase values (the angle of the complex convolution at each time-frequency point) were used to compute two measures of phase consistency i) inter-trial phase clustering (ITPC) and inter-site phase clustering (ISPC) (28). Specifically, the ITPC was computed at each electrode location by measuring the consistency of the instantaneous phase angle values across trials for each time-frequency point. The ISPC was computed as the across-trial consistency of the difference in phase angle values (see SI) for every pair of electrodes within each subject (all combinations: n*(n-1)/2). Electrodes pairs across different arrays (subject N1) or across hemispheres (subject N3) were excluded, leading to 1252 total pairs (ranging from 36 to 465 across subjects). The raw values of both measures (ranging between 0 and 1) were baseline corrected (ranging between −1 and 1) using the average values in the pre-stimulus window (−500 to 0 ms, baseline subtraction) to allow for comparisons and averages across subjects. To perform statistical testing on the ITPC/ISPC time-frequency maps, the measures were transformed (z-fisher) at the group level (one map per subject, with 1 s pre-stimulus and 1 s post-stimulus) and permutation testing was performed to build a null distribution of values (shuffling the pre/post stimulus labels randomly across subjects 1000 times). The time-frequency ITPC/ISPC values were evaluated using alpha level of 0.05 and subsequently a multiple comparison correction was performed by removing clusters of significance smaller than ones occurring by chance (p_corr_<0.05) in the surrogate data.

#### NBG phase synchrony and inter-electrode distance

To evaluate whether ISPC occurring in the NBG range was modulated by the distance between recording sites, we modeled the relation between ISPC values and electrode distances. Specifically, the ISPC values for each electrode-pair (1252 pairs, as defined above) were averaged across time (in the 250-500 ms post-stimulus window) and across frequency (in the NBG range, 20-60 Hz). Inter-electrode distance was defined as the ‘array’ distance between sites (which ranged between 2 mm-8 cm). The relation between the average ISPC values and the distance (mm) between the electrodes forming each pair was modeled by fitting an exponential decay function of the form:

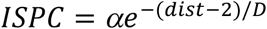

where ISPC is the phase clustering value computed for a pair of electrodes and *dist* is the distance between that pair (in mm, note that the minimum distance between a pair of electrodes is 2 mm and this is subtracted from *dist* to re-center it on 0 mm in the equation). The coefficients α and D were estimated using nonlinear robust minimum absolute deviations fitting (statistics and machine learning toolbox in Matlab). The coefficient α represents the initial value of ISPC at the minimal distance, while D represents the spatial decay constant, thus the distance at which the ISPC decayed to 1/*e* of its initial value. The same model (for details on starting parameters and bounds, see SI) was fitted using three different measures of phase-based synchronization, yielding similar results to ISPC (magnitude squared coherence, phase-lag index and weighted phase-lag index, see SI and Figure S5).

#### Visual grating contrast classification

To further test the relationship between grating contrast levels and induced NBG oscillation frequency, we sought to use NBG and BBG amplitude to classify contrast levels. We employed a support vector machine (SVM) approach using single trial spectral data from the NBG or BBG ranges to build models that would classify the three contrast levels used (20%, 50% & 100%). The model features were vectors of amplitude values for each frequency between 20-60Hz for NBG and 70-150Hz for BBG, obtained by averaging across the 250-500ms window. These vectors of amplitude values were normalized between 0-1 by dividing each value by the maximum amplitude value across the frequency range. Separate models were built for NBG and BBG, with the prediction that NBG would provide higher classification accuracy given the clear changes in peak NBG frequency across contrast levels, compared with BBG. For each electrode, we used the features above to obtain classification accuracy via a leave one out cross validation approach. The SVM model was trained with a linear kernel and the penalty parameter set to 1.

All classification analysis was performed in Matlab (2018a) using functions from the LIBSVM toolbox (https://www.csie.ntu.edu.tw/~cjlin/libsvm/).

Classification accuracy (for stimulus contrast) was calculated across all visually responsive electrodes, separately for NBG and BBG. To test the significance of classification values, we compared observed accuracy with a null distribution created by permutation testing (swapping trial labels prior to SVM training) 1000 times. Therefore, we obtained a classification accuracy and p-value for each electrode, for both NBG and BBG. Finally, as an additional control to confirm classification was only sensitive to responses in early visual cortex, we tested all electrodes sites (beyond occipital lobe) in one subject (N7) as shown in Figure S4. As clearly shown, there was no significant classification of grating contrast levels outside of visual regions.

## Author contributions

B.L.F., W.B. and M.S.B. designed the research; B.L.F., W.B. and D.Y. conducted the experiments; E.B., B.L.F. and Y.L analyzed the data; B.L.F. and E.B wrote the manuscript; all authors edited the manuscript.

## Acknowledgements

We thank Ping Sun for assistance with experimental setup and data recording. This work was supported by NIH grants R01NS065395 to M.S.B; R01EY023336 to D.Y.; R00MH103479 and R01MH116914 to B.L.F.

## References

1. Fries P (2005) A mechanism for cognitive dynamics: neuronal communication through neuronal coherence. Trends Cogn Sci 9(10):474–480.

2. Fries P (2009) Neuronal gamma-band synchronization as a fundamental process in cortical computation. Annu. Rev. Neurosci. 32:209–224.

3. Singer W (1999) Neuronal synchrony: a versatile code for the definition of relations? Neuron 24(1):49–65, 111–125.

4. Gray CM (1999) The temporal correlation hypothesis of visual feature integration: still alive and well. Neuron 24(1):31–47, 111–125.

5. Singer W & Gray CM (1995) Visual feature integration and the temporal correlation hypothesis. Annu. Rev. Neurosci. 18:555–586.

6. Womelsdorf T, et al. (2007) Modulation of neuronal interactions through neuronal synchronization. Science 316(5831):1609–1612.

7. Eckhorn R, et al. (1988) Coherent oscillations: a mechanism of feature linking in the visual cortex? Multiple electrode and correlation analyses in the cat. Biol. Cybern. 60(2):121–130.

8. Gray CM & Singer W (1989) Stimulus-specific neuronal oscillations in orientation columns of cat visual cortex. Proc. Natl. Acad. Sci. U. S. A. 86(5):1698–1702.

9. Fries P, Reynolds JH, Rorie AE, & Desimone R (2001) Modulation of oscillatory neuronal synchronization by selective visual attention. Science 291(5508):1560–1563.

10. Jensen O, Kaiser J, & Lachaux JP (2007) Human gamma-frequency oscillations associated with attention and memory. Trends Neurosci. 30(7):317–324.

11. Yuval-Greenberg S, Tomer O, Keren AS, Nelken I, & Deouell LY (2008) Transient induced gamma-band response in EEG as a manifestation of miniature saccades. Neuron 58(3):429–441.

12. Whitham EM, et al. (2007) Scalp electrical recording during paralysis: quantitative evidence that EEG frequencies above 20 Hz are contaminated by EMG. Clin. Neurophysiol. 118(8):1877–1888.

13. Lachaux JP, Axmacher N, Mormann F, Halgren E, & Crone NE (2012) High-frequency neural activity and human cognition: past, present and possible future of intracranial EEG research. Prog. Neurobiol. 98(3):279–301.

14. Miller KJ, et al. (2014) Broadband changes in the cortical surface potential track activation of functionally diverse neuronal populations. Neuroimage 85 Pt 2:711–720.

15. Ray S & Maunsell JH (2011) Different origins of gamma rhythm and high-gamma activity in macaque visual cortex. PLoS Biol. 9(4):e1000610.

16. Manning JR, Jacobs J, Fried I, & Kahana MJ (2009) Broadband shifts in local field potential power spectra are correlated with single-neuron spiking in humans. J. Neurosci. 29(43):13613–13620.

17. Mukamel R, et al. (2005) Coupling between neuronal firing, field potentials, and FMRI in human auditory cortex. Science 309(5736):951–954.

18. Ray S & Maunsell JH (2010) Differences in gamma frequencies across visual cortex restrict their possible use in computation. Neuron 67(5):885–896.

19. Jia X, Smith MA, & Kohn A (2011) Stimulus selectivity and spatial coherence of gamma components of the local field potential. J. Neurosci. 31(25):9390–9403.

20. Berens P, Keliris GA, Ecker AS, Logothetis NK, & Tolias AS (2008) Comparing the feature selectivity of the gamma-band of the local field potential and the underlying spiking activity in primate visual cortex. Front. Syst. Neurosci. 2:2.

21. Self MW, et al. (2016) The Effects of Context and Attention on Spiking Activity in Human Early Visual Cortex. PLoS Biol. 14(3):e1002420.

22. Hermes D, Miller KJ, Wandell BA, & Winawer J (2015) Stimulus Dependence of Gamma Oscillations in Human Visual Cortex. Cereb. Cortex 25(9):2951–2959.

23. Yoshor D, Bosking WH, Ghose GM, & Maunsell JH (2007) Receptive fields in human visual cortex mapped with surface electrodes. Cereb. Cortex 17(10):2293–2302.

24. Fries P, Scheeringa R, & Oostenveld R (2008) Finding gamma. Neuron 58(3):303–305.

25. Hadjipapas A, Lowet E, Roberts MJ, Peter A, & De Weerd P (2015) Parametric variation of gamma frequency and power with luminance contrast: A comparative study of human MEG and monkey LFP and spike responses. Neuroimage 112:327–340.

26. Cohen AL, et al. (2008) Defining functional areas in individual human brains using resting functional connectivity MRI. Neuroimage 41(1):45–57.

27. Fries P, Nikolic D, & Singer W (2007) The gamma cycle. Trends Neurosci. 30(7):309–316.

28. Cohen MX & Gulbinaite R (2014) Five methodological challenges in cognitive electrophysiology. Neuroimage 85 Pt 2:702–710.

29. Haegens S, Cousijn H, Wallis G, Harrison PJ, & Nobre AC (2014) Inter- and intra-individual variability in alpha peak frequency. Neuroimage 92:46–55.

30. Brunet N, Vinck M, Bosman CA, Singer W, & Fries P (2014) Gamma or no gamma, that is the question. Trends Cogn Sci 18(10):507–509.

31. Hermes D, Miller KJ, Wandell BA, & Winawer J (2015) Gamma oscillations in visual cortex: the stimulus matters. Trends Cogn Sci 19(2):57–58.

32. Hermes D, Miller KJ, Noordmans HJ, Vansteensel MJ, & Ramsey NF (2010) Automated electrocorticographic electrode localization on individually rendered brain surfaces. J. Neurosci. Methods 185(2):293–298.

33. Brunet N, et al. (2015) Visual cortical gamma-band activity during free viewing of natural images. Cereb. Cortex 25(4):918–926.

34. Stigliani A, Weiner KS, & Grill-Spector K (2015) Temporal Processing Capacity in High-Level Visual Cortex Is Domain Specific. J. Neurosci. 35(36):12412–12424.

35. Haller M, et al. (2018) Parameterizing neural power spectra. bioRxiv:299859.

36. Womelsdorf T, et al. (2012) Orientation selectivity and noise correlation in awake monkey area V1 are modulated by the gamma cycle. Proc. Natl. Acad. Sci. U. S. A. 109(11):4302–4307.

37. Bastos AM, et al. (2015) Visual areas exert feedforward and feedback influences through distinct frequency channels. Neuron 85(2):390–401.

38. Bosman CA, et al. (2012) Attentional stimulus selection through selective synchronization between monkey visual areas. Neuron 75(5):875–888.

39. Ray S & Maunsell JH (2015) Do gamma oscillations play a role in cerebral cortex? Trends Cogn Sci 19(2):78–85.

40. Burns SP, Xing D, & Shapley RM (2011) Is gamma-band activity in the local field potential of V1 cortex a “clock” or filtered noise? J. Neurosci. 31(26):9658–9664.

41. Burns SP, Xing D, Shelley MJ, & Shapley RM (2010) Searching for autocoherence in the cortical network with a time-frequency analysis of the local field potential. J. Neurosci. 30(11):4033–4047.

42. Xing D, et al. (2012) Stochastic generation of gamma-band activity in primary visual cortex of awake and anesthetized monkeys. J. Neurosci. 32(40):13873–13880a.

43. Roberts MJ, et al. (2013) Robust gamma coherence between macaque V1 and V2 by dynamic frequency matching. Neuron 78(3):523–536.

44. Nikolic D, Fries P, & Singer W (2013) Gamma oscillations: precise temporal coordination without a metronome. Trends Cogn Sci 17(2):54–55.

45. Kayser C, Salazar RF, & Konig P (2003) Responses to natural scenes in cat V1. J. Neurophysiol. 90(3):1910–1920.

46. Shirhatti V & Ray S (2018) Long-wavelength (reddish) hues induce unusually large gamma oscillations in the primate primary visual cortex. Proc. Natl. Acad. Sci. U. S. A. 115(17):4489–4494.

47. Leszczynski M, et al. (2019) Dissociation of broadband high-frequency activity and neuronal firing in the neocortex. bioRxiv:531368.

48. Buzsaki G, Anastassiou CA, & Koch C (2012) The origin of extracellular fields and currents-EEG, ECoG, LFP and spikes. Nat. Rev. Neurosci. 13(6):407–420.

49. Miller KJ, Zanos S, Fetz EE, den Nijs M, & Ojemann JG (2009) Decoupling the cortical power spectrum reveals real-time representation of individual finger movements in humans. J. Neurosci. 29(10):3132–3137.

50. Winawer J, et al. (2013) Asynchronous broadband signals are the principal source of the BOLD response in human visual cortex. Curr. Biol. 23(13):1145–1153.

51. Lowet E, Roberts MJ, Peter A, Gips B, & De Weerd P (2017) A quantitative theory of gamma synchronization in macaque V1. Elife 6.

52. Logothetis NK, Pauls J, Augath M, Trinath T, & Oeltermann A (2001) Neurophysiological investigation of the basis of the fMRI signal. Nature 412(6843):150–157.

53. Nir Y, et al. (2007) Coupling between neuronal firing rate, gamma LFP, and BOLD fMRI is related to interneuronal correlations. Curr. Biol. 17(15):1275–1285.

54. Foster BL, Rangarajan V, Shirer WR, & Parvizi J (2015) Intrinsic and task-dependent coupling of neuronal population activity in human parietal cortex. Neuron 86(2):578–590.

55. Nir Y, Dinstein I, Malach R, & Heeger DJ (2008) BOLD and spiking activity. Nat. Neurosci. 11(5):523–524; author reply 524.

56. Cardin JA (2016) Snapshots of the Brain in Action: Local Circuit Operations through the Lens of gamma Oscillations. J. Neurosci. 36(41):10496–10504.

57. Sohal VS (2016) How Close Are We to Understanding What (if Anything) gamma Oscillations Do in Cortical Circuits? J. Neurosci. 36(41):10489–10495.

58. Bosking WH, et al. (2017) Saturation in Phosphene Size with Increasing Current Levels Delivered to Human Visual Cortex. J. Neurosci. 37(30):7188–7197.

59. Jenkinson M, Beckmann CF, Behrens TE, Woolrich MW, & Smith SM (2012) Fsl. Neuroimage 62(2):782–790.

60. Cox RW (1996) AFNI: software for analysis and visualization of functional magnetic resonance neuroimages. Comput. Biomed. Res. 29(3):162–173.

61. Dale AM, Fischl B, & Sereno MI (1999) Cortical surface-based analysis. I. Segmentation and surface reconstruction. Neuroimage 9(2):179–194.

62. Groppe DM, et al. (2017) iELVis: An open source MATLAB toolbox for localizing and visualizing human intracranial electrode data. J. Neurosci. Methods 281:40–48.

63. Brainard DH (1997) The Psychophysics Toolbox. Spat. Vis. 10(4):433–436.

64. Henriksson L, Nurminen L, Hyvarinen A, & Vanni S (2008) Spatial frequency tuning in human retinotopic visual areas. J Vis 8(10):5 1–13.

65. R Development Core Team (2010) R: A language and environment for statistical computing (R foundation for Statistical Computing, Vienna, Austria).

