## Supplementary Material for "Distinct Narrow and Broadband Gamma Responses in Human Visual Cortex"

### Supplementary Information Text

### Control Analyses

**Stimulus monitor testing.** We performed repeated measurements of the experimental LCD monitor using a photodiode to ensure the NBG effects reported for Experiment 1 were not confounded by sub-threshold flickering of the stimulus. The power spectrum of photodiode recordings (sampled at 30 kHz) of the stimulus monitor was computed and averaged across three repetitions of experiment 1. The mean power spectrum showed only one peak frequency at 60 Hz, matching the framerate of the LCD monitor. The 60 Hz peak in the power spectrum was not a specific feature of the presentation of the stimuli for experiment 1 (the same monitor was used to present stimuli in experiment 2). In addition, as typically performed, we notched filtered intracranial recordings at 60Hz and harmonics to reduce line noise for both experiments. Therefore, we do not expect that the frame-rate of presentation (or a stimulus flicker) to be a confound on any of the effects reported. Importantly, the group mean NBG peaks in Experiment 1 we observed were 36.4 Hz, 40.2Hz, and 43.8Hz, for the 20%, 50% and 100% contrast conditions respectively (all < 60 Hz; see Figure 1).

**NBG peak frequency modulation.** In the main text, NBG peak frequency for a given electrode was based on identifying the peak frequency at a single-trial level (i.e. the frequency showing the maximal value of amplitude change in the NBG range, 20-60 Hz) and then averaging frequency values across trials (separately for each contrast level). The average values for each electrode were then averaged within subjects (as shown in Figure 2D) and analyzed with a within-subjects ANOVA. We performed two control analyses of the NBG peak frequency and its modulation by grating contrast level to verify that 1) consistent results would be obtained when using a different approach to identify the peak frequency (i.e. using the trial averaged amplitude change spectra) 2) a different analysis would lead to the same modulations (i.e. not averaging across trials and electrodes and including them in a mixed effects analysis).

For the first control analysis, NBG peak frequency for each electrode was estimated by identifying the frequency with maximal value of the trial-averaged normalized amplitude spectra (rather than identifying peak frequencies on single trials). We then averaged the peak frequency values within subjects and analyzed the data following the same approach as in the main text (within-subjects ANOVA). We observed highly similar results (mean peak frequency and standard error for 20% contrast: 33.4±2.4 Hz; 50% contrast: 41.2±2.6 Hz; 100% contrast: 44.7±2.5 Hz; main contrast effect $F(2,12)=19.2$, $p=0.0002$; all three levels of contrast were different from each other, $p<0.05$ Bonferroni corrected). The highly similar results confirm that the modulation of the peak frequency does not vary according to the way the peak is identified (single-trial vs trial-average), suggesting limited variability of NBG peaks across trials within electrodes.

In the second control analysis, we used the single-trial peak NBG frequencies (as in the main text) but we did not average the values across electrodes. We therefore employed a mixed effects model (using lme4 library (1) in R (2)) to evaluate the modulation of the peak NBG frequency, modeling contrast as a fixed effect and electrodes nested in subjects as random effects.

The coefficients showed a similar result as reported in the main text, with a ~3 Hz shift in the peak frequency for each contrast level increment ($\beta_0$, the first fixed parameter captures the 20% contrast level mean: $\beta_0$= 36.8 Hz, s.e. = 1.6 Hz; the other two fixed parameters capture the mean difference between 20% and 50% contrast $\beta_1$ = 3.2 Hz, s.e. = 0.3 Hz; and between 20% and 100% contrast, $\beta_2$ = 6.6 Hz, s.e. = 0.3 Hz). Additionally, to evaluate the significance of the contrast manipulation, the model was compared to a reduced model with the same random effect structure (subjects and electrodes) but no fixed effects (only an intercept term to capture the average peak frequency value, no other parameters to model the mean contrast level differences). The full model had a significantly higher goodness of fit (Akaike information criteria difference with respect to the reduced model = 370; $\chi^2(2)$ = 374.6, p<0.0001) demonstrating that the 2 additional parameters to model the change of peak frequency according to the contrast level significantly increased the amount of variance explained by the model. This confirms that, even when controlling for participant and electrode variation, the contrast of the grating plays a crucial role in explaining the NBG peak frequency value. Overall this analysis validates that the findings reported in the main text were not distorted by averaging across trials and electrode locations within subjects.

**NBG and BBG onset latency.** To validate the reliability of the onset latency values reported in the main text, we used a different algorithm to compute the single-trial onset latency of NBG and BBG amplitude. The first order derivative of the NBG (or BBG) amplitude time series was computed and the median point between the maximum (corresponding to the highest rate of amplitude increase over time) and the neighboring zero-crossing (corresponding to the preceding local minimum) was used as an alternative onset measure (3). This method showed a very high correlation with the method reported in the main text r=0.97, t(131)=39.7, p <0.0001). Both methods were based on selecting the onset features of the amplitude response slope, with the metric reported here being slightly more conservative. Indeed, onset times were estimated to occur ~30 ms later with this alternative method (mean NBG onset = 157 ms, mean BBG onset = 109 ms *vs.* values of 130 ms and 80 ms reported in the main text). Importantly, estimates were simply shifted, as the direction and magnitude of the difference between the average NBG and BBG onset times was unchanged (t(6)=3.8,p<0.01; mean difference ~ 48 ms).

**NBG phase synchrony and inter-electrode distance.** In the main text, we used inter-site phase clustering (ISPC) as a measure of phase-based synchronization, to measure the consistency of the relative NBG phase position recorded at different electrode locations. The ISPC at each time frequency point (t,f) was calculated across trials (from tr = 1 to n) between pairs of electrodes (A and B) as:

$$ISPC_{tf} = \left| \frac{\sum_{tr=1}^{n} e^{i(\omega trA - \omega trB)}}{n} \right|$$

Where $\omega$ is the instantaneous phase angle for trial *tr* at electrode location *A* and *B*, at time *t* and frequency *f*. This measure is also known as the phase-locking statistic or value (PLV, (4); see (5) for nomenclature differences).

Three additional measures were tested to evaluate the generalizability of the result obtained with the ISPC. The first measure we tested was magnitude squared coherence (MSC, (6)), which differs from the ISPC as the phase similarity values are weighted by the power values (cross spectral density of the two signals, normalized by the auto-spectral density of each signal). The second measure was the phase-lag index (PLI, (7)), which differs from the ISPC as the clustering is considered only along the imaginary axis of the complex plane (by taking the sign of the imaginary component). This measure is less influenced by volume-conduction effects (which would have phase angle differences around 0 on the imaginary axis). The third measure was the weighted phase-lag index (wPLI, (8)), which is a modified version of the phase-lag index that weighs more the impact of phase angle differences far from 0 on the imaginary axis, as their sign is more reliable. All measures were baseline corrected (in the same way as for the ISPC, obtaining values ranging from -1 to +1) and used as dependent variable in the nonlinear model (described in the main text). As these measures are sensitive to different attributes of putative synchrony between two regions, they served as controls for our observed relationship between NBG ISPC and inter-electrode distance. We therefore computed the same synchrony-distance relationship for each metric, as shown in Figure S5 B for the 20% contrast level.

The spatial decay constant (D) ranged between ~2 and 4 mm depending on the measure used (20% contrast: MSC-D = 1.8 mm [95% confidence interval: 1.8:1.9]; PLI-D = 3.5 [3.4:3.7]; wPLI-D =3.6 [3.4:3.9]; see Figure S5 for a comparison with ISPC-D: 2.3 [2.2:2.4]). The relationship between synchrony and distance did not seem to be affected by contrast (e.g. 100% contrast: MSC-D = 2.0 [1.9:2.1]; PLI-D = 2.9 [2.6:3.2]; wPLI-D =2.7 [2.2:3.2], ISPC-D = 2.3 [2.2:2.3]), validating the reliability of the result reported in the main text across different metrics.

**Power spectra distribution.** To further test for the occurrence or absence of NBG spectral peaks we adopted an alternative method to parametrize power spectra (9). This method models power spectra as a combination of the 1/frequency component (broadband / aperiodic) in addition to a series of gaussians, which capture the presence of peaks (periodic components). The algorithm developed by Haller et al. first fits an exponential function to model the 1/f relationship (using a robust fit approach, modeling the offset, slope and the presence of an inflexion point) and subsequently removes the fitted component, obtaining a flattened spectrum. Next, the flattened spectrum is convolved with a series of gaussians to model the presence/absence of peaks in the spectrum that were not accounted for by the aperiodic component. The presence of an oscillation or, more generally, of a peak in the spectrum will result in a successful fit of the gaussian model, capturing the center frequency, amplitude, and bandwidth of the peak. After these steps, the full model is built based on a combination of the parameters obtained from the aperiodic fit and gaussian fits.

We applied this method to our average power spectra values (group-level averaged, obtained by considering the 250-500 ms post stimulus window for the task data, and 200 ms pre-stimulus baseline for comparison: Figure 5). The model was fit on the frequency range between 20 and 150 Hz, by allowing the aperiodic component to have an inflexion point and by constraining the gaussian fits to have a bandwidth between 2 and 80 Hz and to be at least 3 standard deviations above the aperiodic signal. When considering the grating task, all the contrast levels exhibited peaks with values similar to those obtained by our peak analysis in the main text (20% contrast = 31 Hz; 50% contrast = 36.5 Hz; 100% contrast = 41.2 Hz; ~9 Hz bandwidth). In opposition, the power spectra for the natural image category task resulted in model fits with no identifiable periodic components (for all categories). The aperiodic component parameters were similar across categories (offset between 11-11.6, slope between 3.6 and 3.8, inflexion point occurring between 24 and 31 Hz). The baseline power for both tasks did not exhibit any peaks and it required a slightly different model for the aperiodic component, as the inflexion point was not present. The offset and slope parameters were highly similar for the two baselines (offset: 10.6-10.7, slope 3.7 for both).

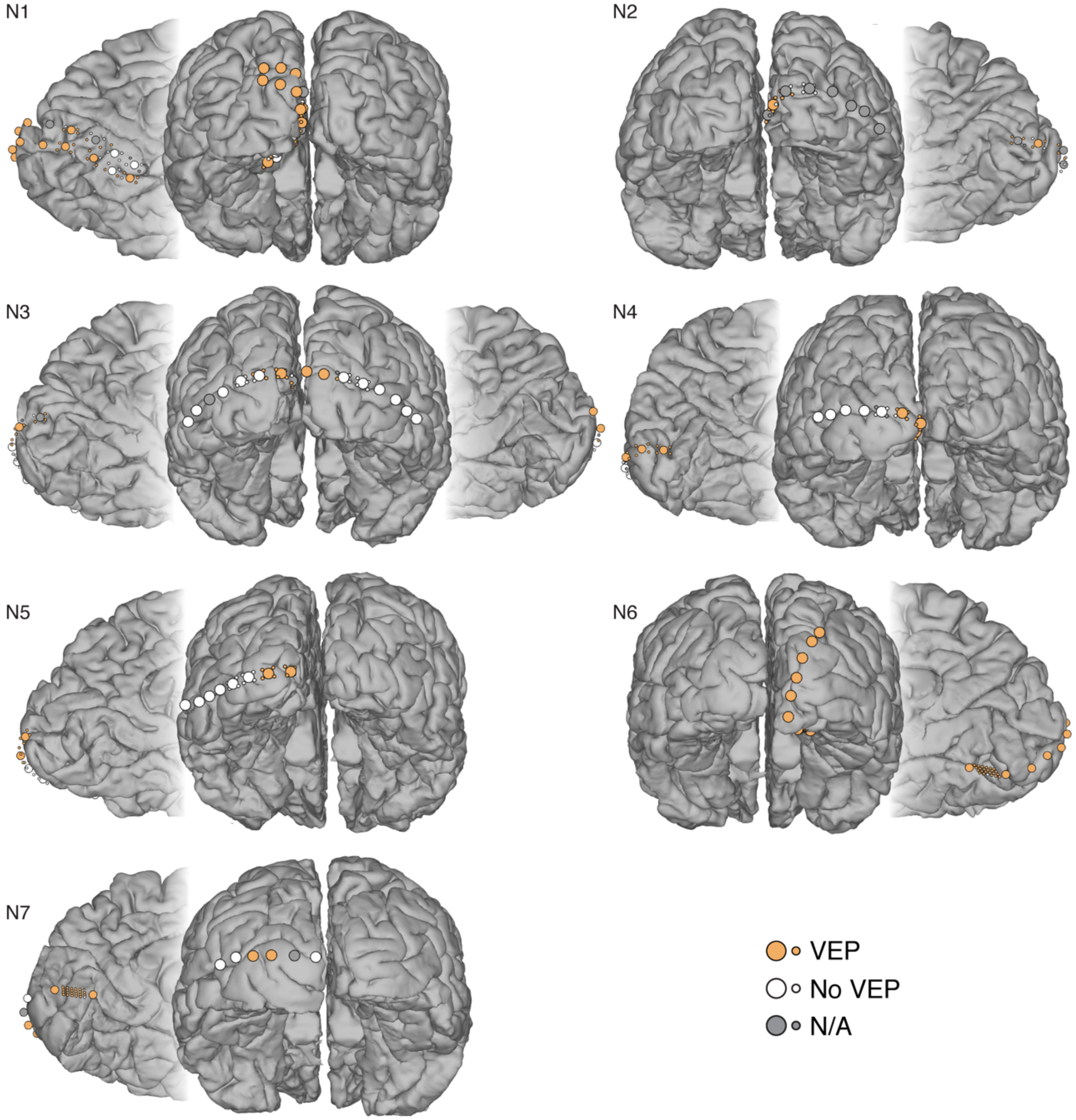


**Fig. S1. Single subject cortical surface and electrode array location**. Electrodes are color-coded in a similar fashion to Figure 1: orange indicates electrodes exhibiting a VEP, white indicates electrodes that did not. Additionally, grey indicates electrodes that were excluded from data analysis (Electrode type A for subjects N1-5 and type B for N6-7, as described in the Methods and schematized in Figure 1 & Figure S5).

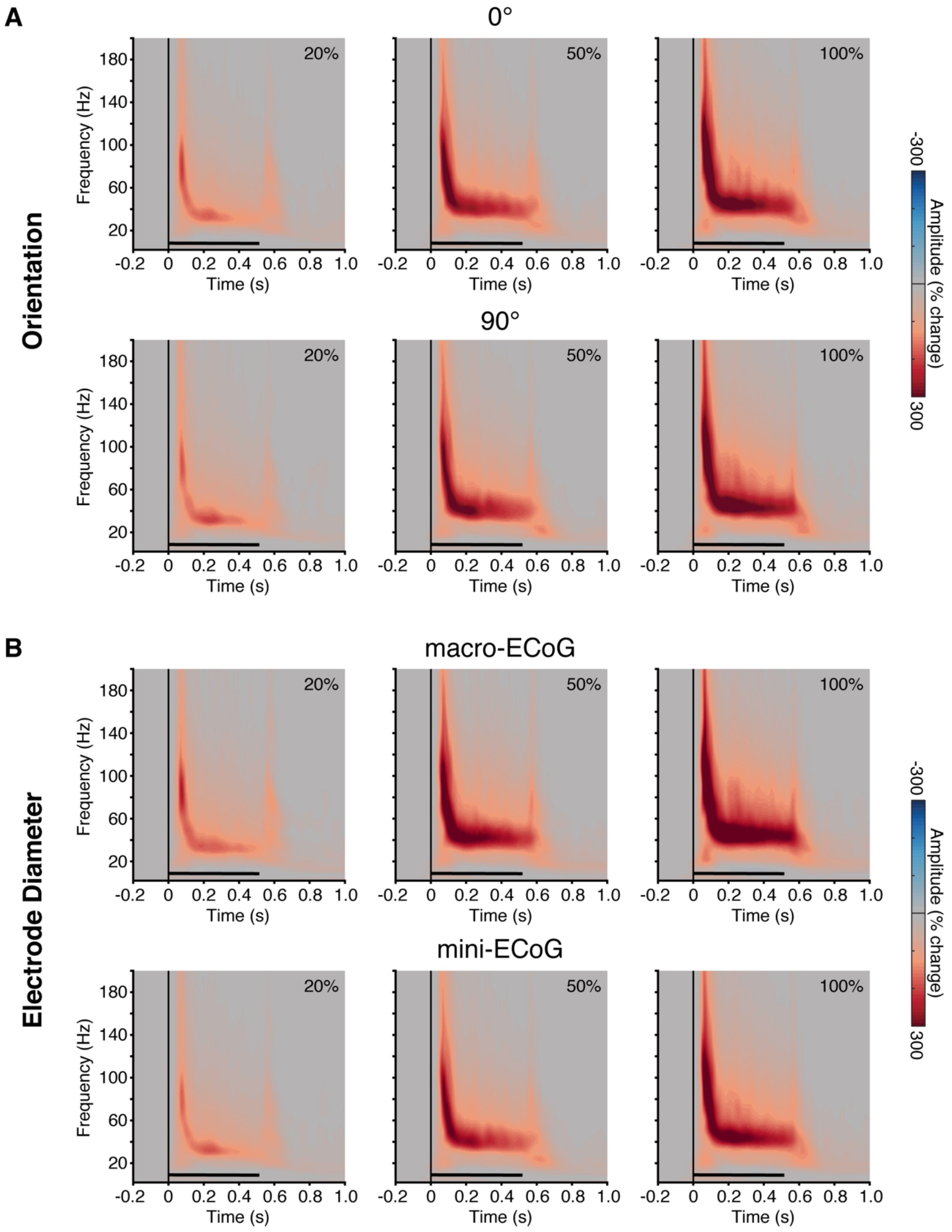


**Fig. S2. Influence of grating orientation and electrode size on spectral response.** Spectrograms show group average time-frequency responses for the 20%, 50% and 100% contrast levels. **A)** Spectrograms for the two grating orientations (0° and 90°). **B)** Spectrograms for the two electrode sizes (macro- and mini-ECoG). As clear from both panels, the observed spectral responses show a striking similarity to Figure 1, supporting the combination of orientation and electrode diameter in data analyses.

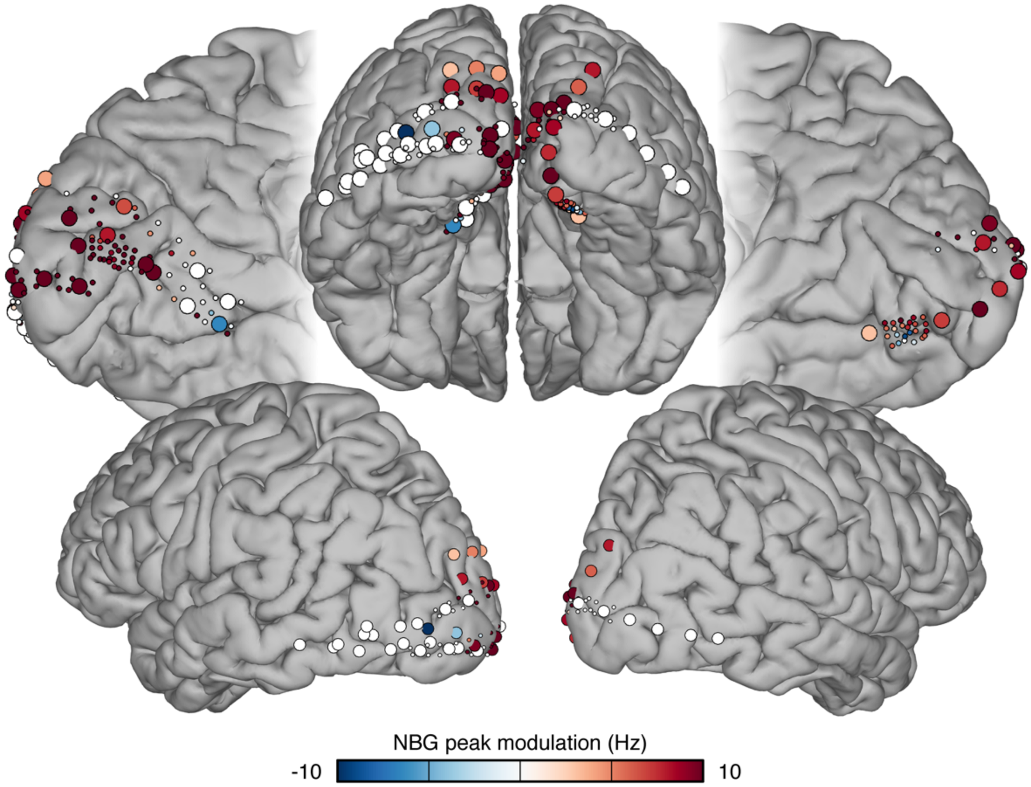


**Fig. S3. Magnitude of shift in NBG peak frequency**. The contrast modulation of NBG peak frequency is represented at each electrode location (on the standard brain shown in Figure 1) and color coded according to the magnitude of frequency shift (by using the difference between the average peak frequency at 100% contrast versus 20% contrast). Note that few electrodes showed an apparent opposite modulation of the peak frequency: these electrodes did not display clear spectral increases in the lower contrast level causing a noisy estimate of the peak frequency in the NBG range.

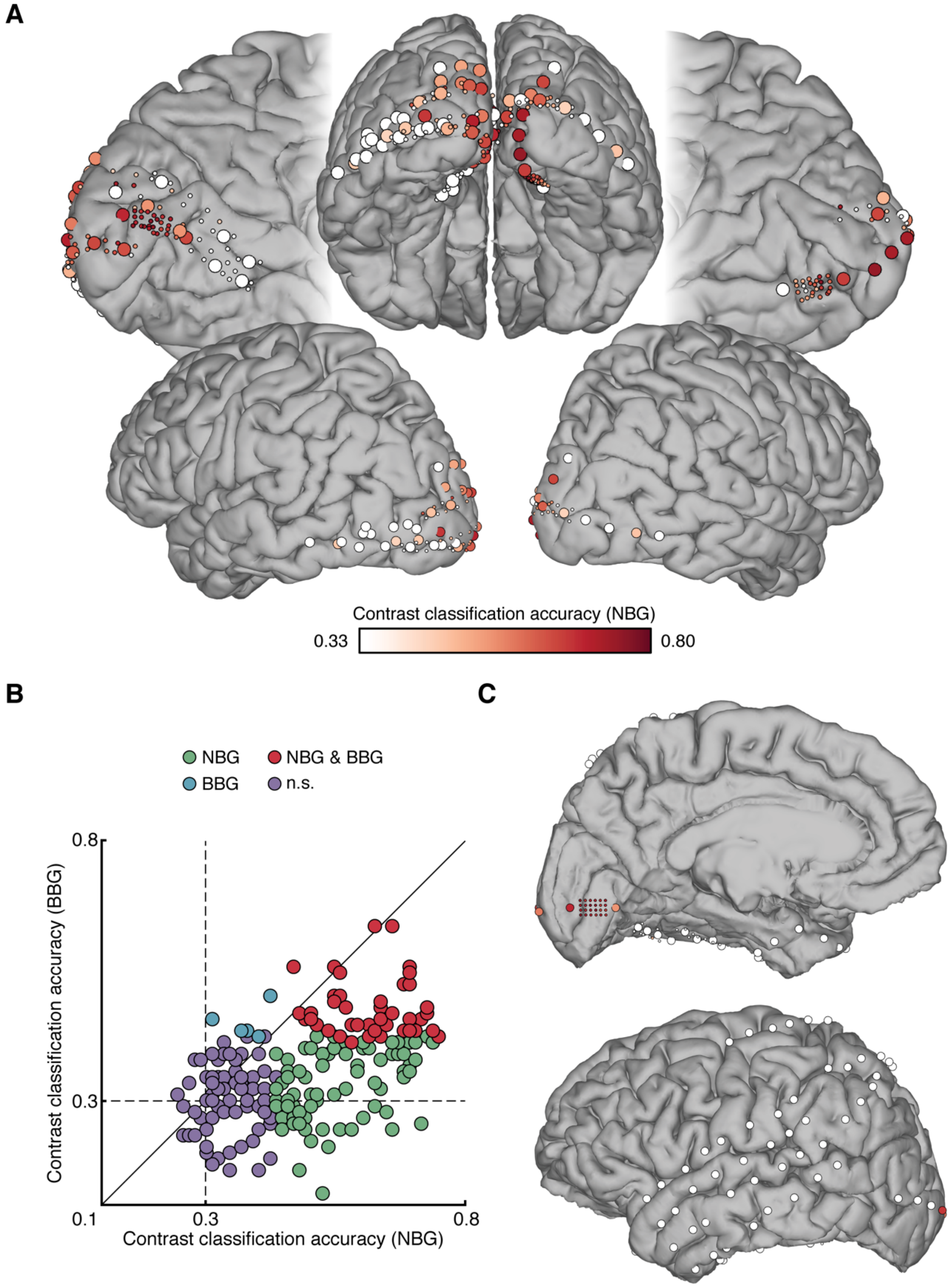


**Fig. S4. Classification of grating stimulus contrast. A)** Group data showing grating contrast level classification accuracy for each electrode (on the standard brain shown in Figure 1) based on a support vector machine (SVM) using averaged NBG amplitude (see Methods). Electrode locations with non-significant classification accuracy (see Methods) are shown in white. Note that all occipital electrodes were included in this analysis, validating the selection of the electrodes based on the presence of the VEP as a criterion that was able to capture responsive electrodes. There was ~80% agreement between electrodes with above chance classification accuracy and those showing a VEP (see Figure 1 for a visual comparison). **B)** Scatter plot shows classification accuracy for the NBG-trained SVM plotted against the classification accuracy for the BBG-trained SVM for each electrode. Values are color coded according to the statistical significance of their classification accuracy (assessed with permutation testing, see Methods). **C)** Same as panel A for an individual subject (N7) showing that no electrode locations beyond the occipital lobe had above chance classification accuracy (using NBG; color map same as A).

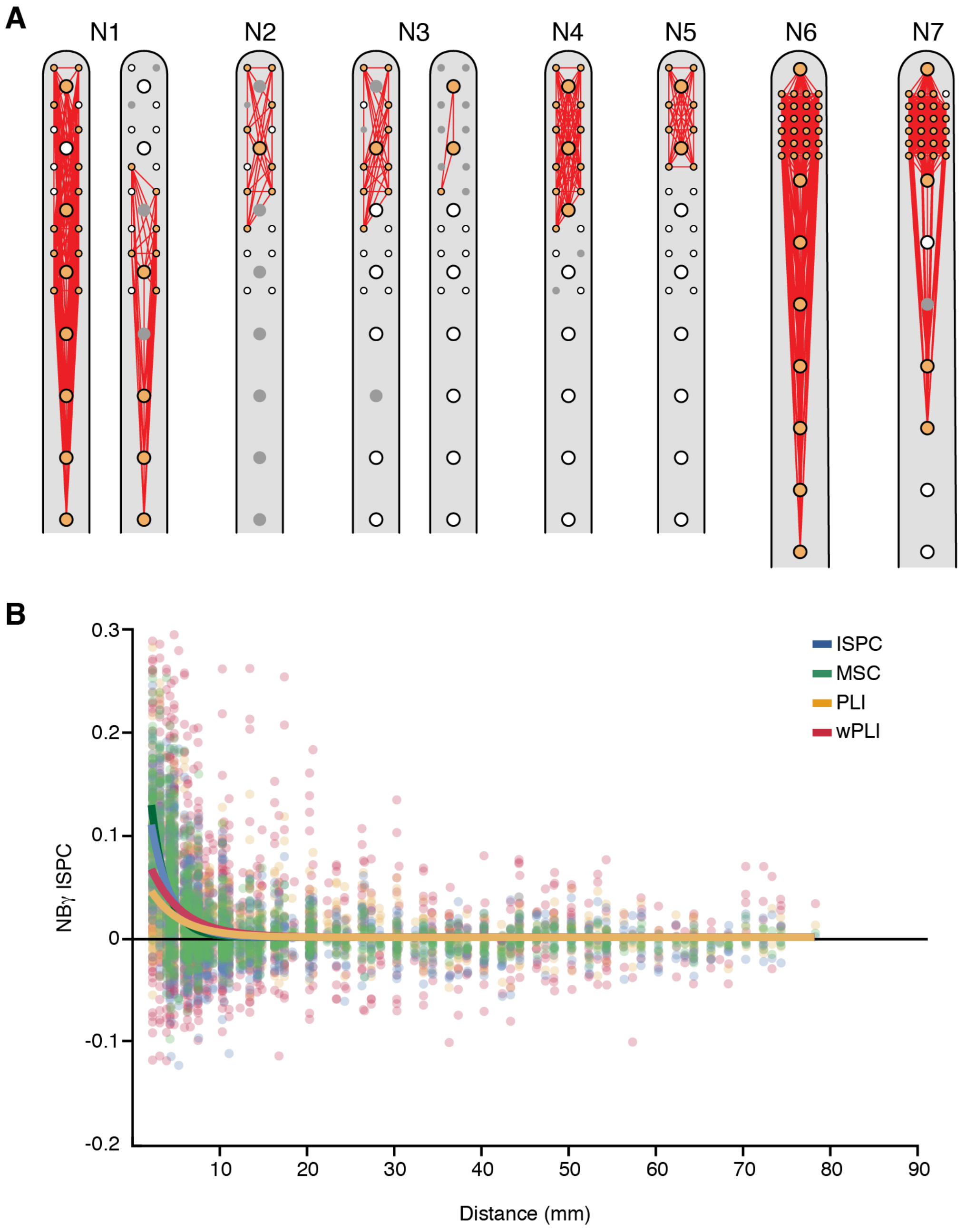


**Fig. S5. Electrode pairs and control metrics for NBG synchrony analysis. A)** Electrode array configurations for all subjects (N1-7), where red lines indicate all electrode pairs used in the phase-based synchrony analyses (electrode colors same as Figure S1). **B)** Scatter plot shows a comparison between the decay of NBG phase based synchrony over inter-electrode distance using different metrics (ISPC, as in main text, MSC: mean squared coherence, PLI: phase lag index and wPLI: weighted phase lag index). Data is shown for the 20% contrast condition.

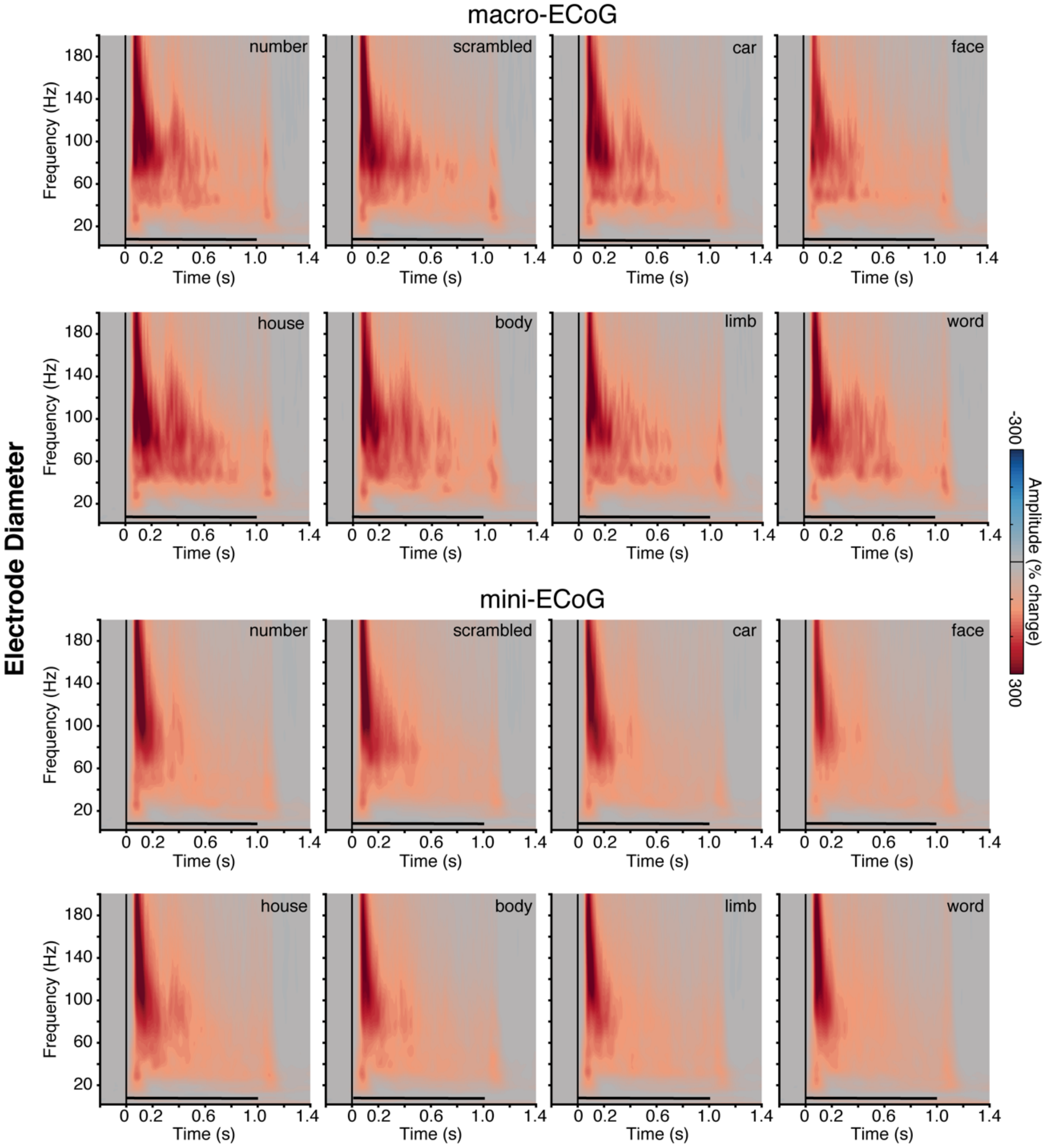


**Fig. S6. Influence of electrode size on spectral response (Experiment 2).** Spectrograms show group average time-frequency responses for the natural image categories in experiment 2. Spectrograms for the two electrode sizes (macro- and mini-ECoG) display similar responses to those reported in Figure 4, where electrode size was collapsed in data analyses. Macro-ECoG responses show some qualitative differences, with a longer duration BBG response, however remains highly distinct from responses observed for experiment 1.

**Table S1. Subject information.** Table shows subject sex, age at time of experiment, implant hemisphere and electrode array type (A or B), electrode count in the region of interest (ROI) and the electrode count selected by the presence of a visual evoked potential (VEP).

| Subject number | Sex | Age | Electrode array Configuration | | #Electrodes in ROI | #Electrodes showing VEP |
|---|---|---|---|---|---|---|
| | | | Left hemisphere | Right hemisphere | | |
| **N1** | M | 32 | A(x2) | - | 44 | 29 |
| **N2** | M | 44 | - | A | 16 | 9 |
| **N3** | M | 54 | A | A | 36 | 13 |
| **N4** | M | 20 | A | - | 22 | 14 |
| **N5** | M | 47 | A | - | 24 | 10 |
| **N6** | F | 37 | - | B | 32 | 31 |
| **N7** | M | 25 | B | - | 31 | 27 |

**References**


1. Bates D, Mächler M, Bolker B, & Walker S (2014) Fitting linear mixed-effects models using lme4. *arXiv preprint arXiv:1406.5823*.
2. R Development Core Team (2010) R: A language and environment for statistical computing (R foundation for Statistical Computing, Vienna, Austria).
3. Bartoli E, Aron AR, & Tandon N (2018) Topography and timing of activity in right inferior frontal cortex and anterior insula for stopping movement. *Hum. Brain Mapp.* 39(1):189-203.
4. Lachaux JP, Rodriguez E, Martinerie J, & Varela FJ (1999) Measuring phase synchrony in brain signals. *Hum. Brain Mapp.* 8(4):194-208.
5. Cohen MX (2014) *Analyzing neural time series data: theory and practice* (MIT press).
6. Shils JL, Litt M, Skolnick BE, & Stecker MM (1996) Bispectral analysis of visual interactions in humans. *Electroencephalogr. Clin. Neurophysiol.* 98(2):113-125.
7. Stam CJ, Nolte G, & Daffertshofer A (2007) Phase lag index: assessment of functional connectivity from multi channel EEG and MEG with diminished bias from common sources. *Hum. Brain Mapp.* 28(11):1178-1193.
8. Vinck M, Oostenveld R, van Wingerden M, Battaglia F, & Pennartz CM (2011) An improved index of phase-synchronization for electrophysiological data in the presence of volume-conduction, noise and sample-size bias. *Neuroimage* 55(4):1548-1565.
9. Haller M*, et al.* (2018) Parameterizing neural power spectra. *bioRxiv*:299859.
